# A SARS-CoV-2 – host proximity interactome

**DOI:** 10.1101/2020.09.03.282103

**Authors:** Payman Samavarchi-Tehrani, Hala Abdouni, James D.R. Knight, Audrey Astori, Reuben Samson, Zhen-Yuan Lin, Dae-Kyum Kim, Jennifer J. Knapp, Jonathan St-Germain, Christopher D. Go, Brett Larsen, Cassandra J. Wong, Patricia Cassonnet, Caroline Demeret, Yves Jacob, Frederick P. Roth, Brian Raught, Anne-Claude Gingras

**Author notes:** equal contribution (Hala Abdouni, James D.R. Knight).

## Abstract

Viral replication is dependent on interactions between viral polypeptides and host proteins. Identifying virus-host protein interactions can thus uncover unique opportunities for interfering with the virus life cycle via novel drug compounds or drug repurposing. Importantly, many viral-host protein interactions take place at intracellular membranes and poorly soluble organelles, which are difficult to profile using classical biochemical purification approaches. Applying proximity-dependent biotinylation (BioID) with the fast-acting miniTurbo enzyme to 27 SARS-CoV-2 proteins in a lung adenocarcinoma cell line (A549), we detected 7810 proximity interactions (7382 of which are new for SARS-CoV-2) with 2242 host proteins (results available at covid19interactome.org). These results complement and dramatically expand upon recent affinity purification-based studies identifying stable host-virus protein complexes, and offer an unparalleled view of membrane-associated processes critical for viral production. Host cell organellar markers were also subjected to BioID in parallel, allowing us to propose modes of action for several viral proteins in the context of host proteome remodelling. In summary, our dataset identifies numerous high confidence proximity partners for SARS-CoV-2 viral proteins, and describes potential mechanisms for their effects on specific host cell functions.

## Introduction

SARS-CoV-2 is a positive-sense single stranded RNA (+ssRNA) coronavirus, responsible for the COVID-19 pandemic^1^. No specific therapeutics are currently available for this novel virus. A deeper understanding of SARS-CoV-2 pathobiology at the molecular level will both provide critical insight into the strategies used by the virus to subvert and hijack host cell functions, and identify virus-host protein-protein interactions that are required for viral replication, many of which could represent useful drug targets.

The SARS-CoV-2 genome consists of a capped and polyadenylated RNA of approximately 30kb, containing one large 5’ open reading frame (ORF), which encodes two large “polyproteins”. The polyproteins are proteolytically processed by viral proteases to yield the non-structural proteins NSP1-NSP16. 13 additional smaller ORFs (n.b. whether ORF10 is expressed as a protein remains to be demonstrated^2^) encode the structural proteins – spike (S), envelope (E), membrane (M) and nucleocapsid (N) – and other polypeptides. Together, these proteins comprise the SARS-CoV-2 proteome^3^.

RNA (+) viruses such as SARS-CoV-2 effect a remarkable re-organization of host cell ER-Golgi membranes to form single membrane invaginations and double membrane vesicles (DMV), which serve to organize and protect the viral replication machinery, and which shield new virus particles from innate immune detection^4,5^. Virus-host interactions that take place in replication organelles are obvious targets for therapeutic intervention, yet these membrane protein interactions have remained poorly characterized.

Three recent studies utilized affinity purification combined with mass spectrometry (AP-MS) to map over 1000 putative SARS-CoV-2 virus-host protein-protein interactions^6–8^ in HEK293T and A549 cells. While these traditional affinity purification approaches have been of immense value in characterizing biochemically stable protein complexes, this approach is less amenable to the identification of weak or transient interactions, and is not optimal for the identification of interactions that take place at poorly soluble intracellular locations, such as membranes or on chromatin^9^. Of the 29 SARS-CoV-2 proteins, nine possess one or more predicted transmembrane domains. Relatively few interaction partners have been identified for this group of proteins to date (including in the aforementioned studies), suggesting that complementary interaction identification approaches could provide important additional insight into the virus-host interactome.

Proximity-dependent biotinylation approaches are well-suited for defining relationships between proteins in living cells. The BioID approach, which utilizes an abortive bacterial biotin ligase to mediate proximity-dependent biotinylation of proteins in the vicinity of a bait^10^, has been particularly useful for mapping poorly soluble intracellular structures such as membranes^11,12^ and membraneless^13,14^ organelles, and for characterizing the proximal interactomes of other virus proteins^15^. In this approach, a cloud of activated biotin (with a ~5-10nm radius) is generated around the “bait” protein, resulting in the covalent biotinylation of lysine moieties in nearby “prey” proteins. Covalent biotinylation enables the use of harsh lysis conditions to solubilize all structural elements in the cell, including organelles, protein complexes and membranes, and permits the identification of biotinylated proteins using streptavidin affinity purification followed by mass spectrometry. The original BioID was performed with an *E. coli* BirA enzyme harboring a single mutation (R118G), and which displays relatively low catalytic activity (reviewed in ^16^). The application of directed molecular evolution yielded new BirA variants such as miniTurbo, an N-terminally truncated BirA protein with a dozen additional mutations, and dramatically improved catalytic activity^17^. miniTurbo enables efficient biotinylation within minutes, rather than the many hours required for the “classic” BirA* enzyme. Also of critical importance, using our lentivirus-based toolkit (originally established for BirA*^18^), miniTurbo-tagged bait proteins can be expressed in many different cell types.

Here, using our lentiviral delivery system, we conduct miniTurbo BioID on 27 SARS-CoV-2 proteins, tagged at both the N- and C-termini, characterizing their intracellular localization, and mapping virus-host proximity interactomes in the A549 human lung alveolar epithelial cell line. We also include 17 host proteins previously used as subcellular compartment markers, allowing us to report more precisely on the location of the viral proteins, and in some cases propose mechanisms of action.

## Results

### Proximity interactome of SARS-CoV-2 proteins in A549 cells

To generate a proximity interaction map of the SARS-CoV-2 proteome, 27 viral ORFs (as defined in ^19^; Fig. 1a; Supplementary Table 1) were cloned into lentiviral expression vectors^18^, enabling tetracycline-inducible expression of polypeptides fused in-frame with the miniTurbo enzyme^17^ at their amino (_Nt) or carboxy (_Ct) termini. A549 lung adenocarcinoma cells were transduced with lentivirus (see Methods for details on optimization and validation), treated with doxycycline for 24 hours to induce transgene expression, then supplemented with biotin for 15 minutes to initiate proximity-dependent biotinylation. Cells were harvested using stringent lysis conditions for immunoblotting or BioID, or fixed for immunofluorescence. Bait expression, localization and induction of biotinylation were assessed by immunoblotting and immunofluorescence (see below).

**Fig. 1:**
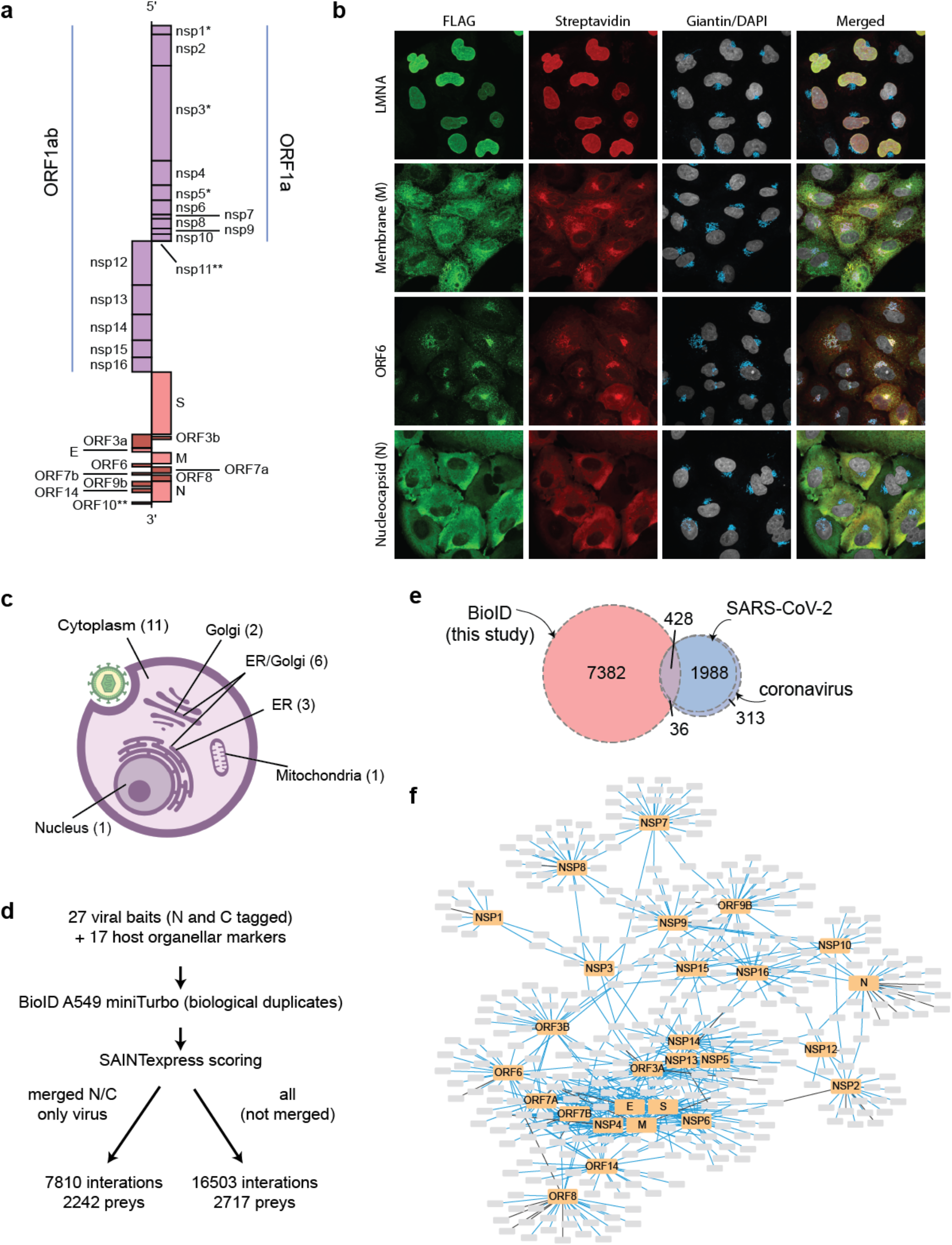
Generation of a SARS-CoV-2 – host proximity interactome in A549 cells. **a**, Schematic of the virus proteins profiled in this study, mapped on the genomic viral RNA (*profiled in wild type and mutant forms; **not profiled). **b**, Representative immunofluorescence images demonstrating miniTurbo-tagged SARS-CoV-2 proteins targeting to different subcellular compartments and miniTurbo-tagged LMNA as a representative host protein (also see Extended data Fig. 2a and b). **c**, Summary of immunofluorescence data (number of baits in each category indicated in parentheses; see Supplementary Table 1 and Extended data Fig. 2a and b). **d**, Overview of the dataset. **e**, Overlap of the proximity interactions reported here with previously reported interactions for SARS-CoV-2 or any coronavirus (see Supplementary Table 3 and Extended Data Fig. 3). **f**, Cytoscape representation of the most abundant (length normalized spectral counts; up to 25 are shown) proximity interactors for each viral bait (See Supplementary Table 4 and covid19interactome.org/#network for an interactive version of this image).

A pilot immunofluorescence study (using transient transfection in HEK293T cells; data not shown) indicated that several SARS-CoV-2 bait proteins localized generally to cytoplasm, but many proteins were instead clearly associated with the endoplasmic reticulum (ER), Golgi apparatus, mitochondria, and possibly other intracellular membranes. Leveraging our previous study on the subcellular organization of a human cell^20^ (in which 192 subcellular markers were profiled by BioID in HEK293 cells, using the classic BirA* enzyme), 17 ER, Golgi, mitochondria and other organellar membrane and non-membrane “marker” ORFs were also cloned into the miniTurbo lentiviral vector, and BioID conducted as above. To model background and non-specific biotinylation, we used three different negative controls: an EGFP vector (not expressing miniTurbo), and miniTurbo fused to either EGFP or a Nuclear Export Sequence (NES).

The majority of the SARS-CoV-2 bait proteins were successfully expressed in A549 cells in both N- and C-terminal tagged configurations (Extended data Fig. 1), and all but NSP1 and NSP3 (which we profiled instead by transient transfection; see Methods) were well expressed in at least one orientation. For NSP1, this observation is consistent with a previously characterized strong suppression of host translation by the SARS-CoV ortholog^21^. A double point mutant defined in SARS-CoV NSP1 that prevents translational inhibition^22^ was introduced in SARS-CoV-2 NSP1 (K164A and H165A). These amino acid changes allowed for the successful production of lentivirus, and expression of the miniTurbo fusion gene in A549 cells. Catalytically inactive mutants of the proteases NSP3 (C857A) and NSP5 (C145A) were also created via mutation of active site residues, as previously described^6^. Together, our dataset thus comprises the proximity interactomes of 27 WT virus baits, mutant versions of NSP1, NSP3 and NSP5, and 17 host baits (**Supplementary Table 1)**.

In addition to profiling the proximity interactome of each of the SARS-CoV-2 proteins, we assessed their subcellular localization using immunofluorescence microscopy in A549 cells. This revealed the localization of 11 baits to the endoplasmic reticulum (NSP4, E, ORF8), the Golgi (ORF3B, ORF7A) or both compartments (NSP6, S, ORF3A, M, ORF6, ORF7B); Fig.1b-c; Extended data Fig. 2a,b; Supplementary Table 1). NSP2 and Nucleocapsid (N) exhibited nuclear exclusion, ORF9B localized to mitochondria localized to the nucleus. NSPs 7-16 exhibited non-discrete localization, distributing throughout the cell. We note that some baits (including ORF3A, ORF7A, ORF14 and ORF8) were detected in different compartments depending on tag location. This is not completely surprising since ORF3A is a multipass membrane protein and ORF7A and ORF8 contain an N-terminal signal sequence. Overall, our results agree to a large extent with previous reports of the subcellular localization of these proteins, and suggest that the miniTurbo tag does not dramatically affect localization.

All viral baits, host subcellular markers and negative controls were profiled (≥2 biological replicates) by BioID and mass spectrometry (see Methods). The dataset was analyzed using Significance Analysis of INTeractome (SAINTexpress^23^), identifying 7810 high-confidence (i.e. Bayesian FDR ≤1%) proximity interactions (after merging N and C-terminal data) for the WT viral baits, with 2242 unique human proteins (Fig. 1d; Supplementary Table 2). The viral proteins recovered between 6 (NSP12) and 938 (M) proximity interactions, with a median of 205, consistent with the localization of many of these proteins to membranous compartments, which tend to produce rich BioID profiles^20^. Across the entire dataset, including host subcellular markers (and without merging the viral bait orientations), 16503 high-confidence proximity interactions with 2715 unique prey proteins were uncovered.

### Comparison to other datasets

We next compared our BioID results with those of recently reported AP-MS results^6–8^, as annotated in BioGRID^24^. A relatively high proportion (21.7%) of previously reported SARS-CoV-2 AP-MS interactions was recapitulated in our study (Fig. 1e, Supplementary Table 3). However, this value varied widely between baits, ranging from 0 (e.g. NSP14) to 43.9% (for ORF8) of previously reported interactions recovered (Extended data Fig. 3). At least 10% of previously reported interactions were recovered for 13 different virus proteins. This group of interactions, cross-validated in multiple approaches, may represent promising therapeutic targets (specific examples discussed below). As expected, BioID expands upon the previously reported AP-MS interactions (1976) to include 7810 proximal interactions, which encompass both direct interactors and proteins residing in the vicinity of the bait protein. We note that 94.5% of our interactions are novel for SARS-CoV-2 (94.1% when expanding to the complement of all coronavirus protein orthologs – SARS-CoV, SARS-CoV-2, MERS – annotated in BioGRID), providing opportunities to gain new insight into the molecular mechanisms by which SARS-CoV-2 proteins subvert and hijack cellular functions.

### Website

To make this resource readily available to the scientific community, we created the covid19interactome.org, a website that can be used to explore the virus-host proximity interactome dataset. Users can explore the website from the point of view of a viral protein with a detailed report on proximity interactions for the WT or mutant variants, or by searching for a specific host prey protein, which will retrieve baits that identify the prey and offer a heat map profile view across the data set. A network exploration tool is also available that displays up to 25 of the most abundant preys (length normalized spectral counts) for each bait (Fig. 1f; Supplementary Table 4). Examples of the functionality of the website are shown in Extended data Fig. 4–6.

### BioID enrichment and profile similarity between viral and host baits

As expected, BioID interactomes for each viral protein were enriched for Gene Ontology (GO) Cellular Compartment (CC) terms matching their immunofluorescence localization. Baits with more diffuse staining or low recovery of proximity interactors did not display any well-defined enrichment for a specific compartment (Fig. 2a and Supplementary Table 5; also see domain and region enrichments within this table). To more precisely define the localization of the viral proteins, both viral and host baits were subjected to clustering by pairwise Jaccard similarity (Fig. 2b). This largely clustered viral baits with host bait proteins with known subcellular location and function, and whose BioID profiles serve as compartment reference signatures. These results also supported the immunofluorescence localization and the enrichment of specific GO terms.

**Fig. 2:**
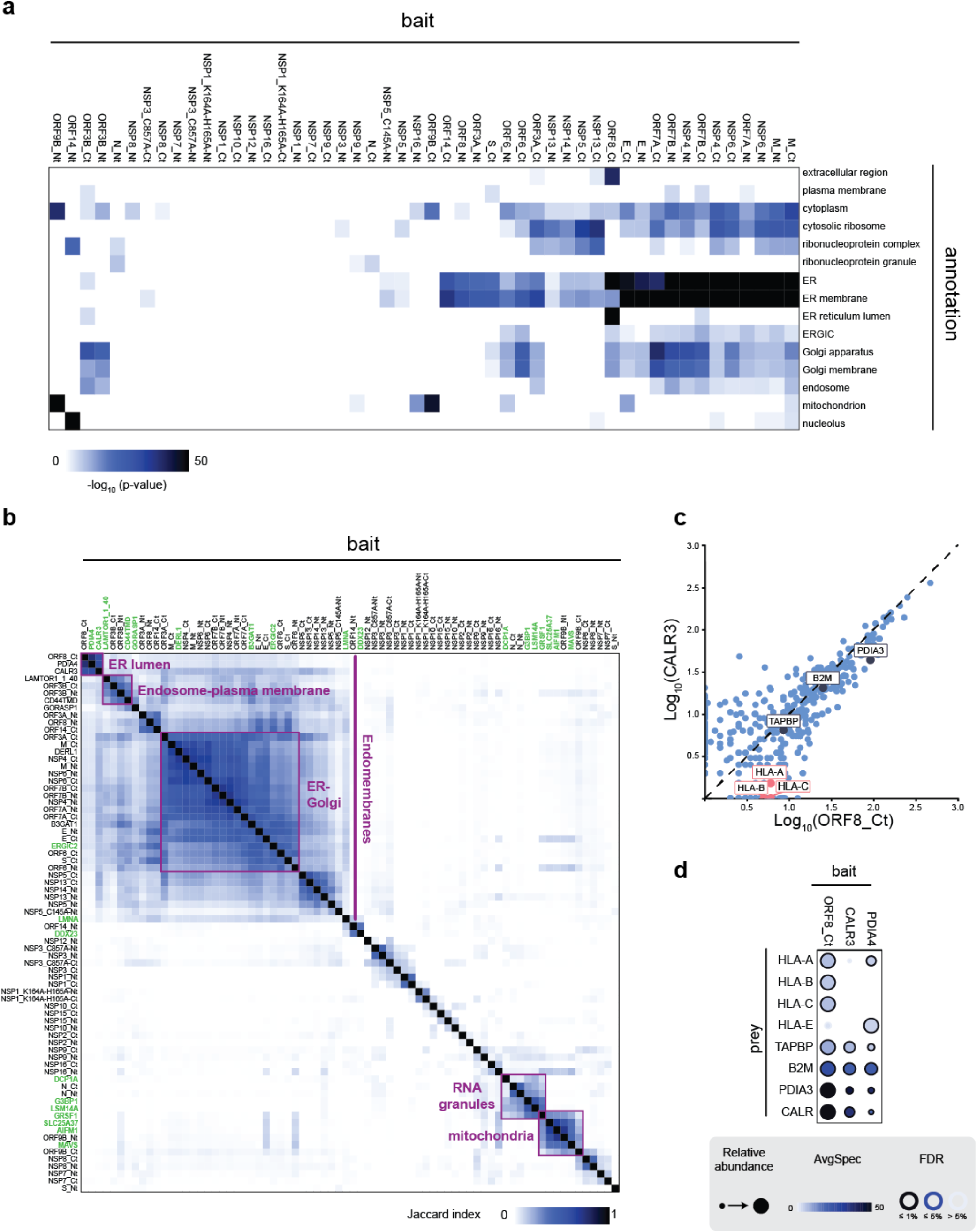
A global proximity interactome view of subcellular localization of SARS-CoV-2 baits. **a**, Enriched GO terms were selected from representative compartments (see Supplementary Table 5 for full list of terms). Baits were clustered using the Euclidean distance and the Ward clustering metric. The −log10 of the adjusted p-value was capped at 50 for display purposes. **b**, The Jaccard index was calculated between each pair of baits using their preys. Baits were clustered using the Euclidean distance and the single clustering metric. Host baits are color coded in green. **c**, Bait-bait comparison (spectral counts) between ORF8_Ct and the ER marker CALR3. Components of the MHC-I complex are marked. **d**, Dotplot of selected ORF8_Ct interactors associated with MHC-I.

A large, dense cluster of host and viral baits localized to the ER and Golgi compartments was clearly identified, defined by host baits that label the cytoplasmic face of these structures: DERL1 (Derlin-1; ER), B3GAT1 (Beta-1,3-Glucuronyltransferase 1, Golgi) and ERGIC2 (ER-Golgi intermediate compartment 2). Within this cluster, viral proteins M, NSP4, NSP6, ORF7A and ORF7B (both N- and C-tagged) most resembled DERL1, while E and ORF6 (both N- and C-tagged) more closely resembled the Golgi and ERGIC markers. Extension of this cluster to include the region marked by host baits localized to lysosomal and plasma membrane/recycling endosome systems (fragments of LAMTOR1 and CD44; see Supplementary Table 1) revealed similarity to ORF3B. Additional baits (ORF3A, NSP5, NSP13) were associated with the larger endomembrane system.

Smaller clusters containing both host and viral proteins were also identified, including: the mitochondrial host baits (GRSF1, SLC25A37, AIFM1, MAVS) and SARS-CoV-2 ORF9B; and the RNA granule host baits (DCP1A, G3BP1, LSM14A, GRSF1) and the viral N protein (both discussed further below).

In most instances, interactome data generated by viral proteins tagged at the N- and C- termini clustered closely with one another, suggesting that both bait variants sample similar environments (as would be expected for non-membrane-spanning soluble proteins or multipass proteins with both termini facing the same side of a given membrane). This includes the aforementioned components of the ER-Golgi-endosome, mitochondria, and RNA granule clusters, but also pairs of viral proteins not associated with any host clusters, and in general recovering fewer proximity partners (i.e. NSP1, NSP2, NSP3, NSP7, NSP8, NSP9 and NSP16).

While similarity in profiles between N and C-terminally tagged variants seemed to be the rule here, there were a few exceptions. First, some of the baits (S, NSP12, NSP13 and NSP14) were clearly better expressed (Extended Data Fig. 1) and recovered meaningful data in only one tagging orientation (e.g. S_Ct clustered with the Golgi-ERGIC and NSP12_Nt and NSP14_Nt were in the larger membrane cluster while only 0–1 confident proximity partners were recovered in the reverse orientation, precluding meaningful clustering). However, two viral proteins that were well expressed in both tag orientations displayed strikingly different profiles depending on tag location. When C-terminally tagged, ORF14 predominantly enriched ER components, while the N-terminally tagged form strongly enriched nucleolar and nuclear proteins (Fig. 2a), consistent with their respective immunofluorescence results (Extended Data Fig. 2). This protein (also known as ORF9c^6^), was recently shown to associate with a variety of membranes when N-terminally tagged with two Strep-tag II peptides^25^, which is in contrast to our results for the N-terminal tag variant. Whether the fusion of the larger miniTurbo moiety to the N-terminus of ORF14 disrupts its localization, or (at least some of) this protein is also localized to the nucleolus remains to be determined. The other notable exception was ORF8, whose C-terminal fusion clustered with the ER lumen proteins CALR3 (Calreticulin-3) and PDIA4 (Protein disulfide-isomerase A4). Interestingly, ORF8_Ct was the only viral protein found to cluster with the ER lumen marker baits. These results are consistent with the fact that ORF8 possesses an N-terminal signal sequence (which would be masked by N-terminal tagging). The overlap between the proximity partners of ORF8_Ct and the two ER lumen markers was extensive (see comparison with CALR3 in Fig. 2c), though there were some proteins, especially in the lower abundance range, that seemed preferentially recovered with ORF8. Interestingly, these included components of the major histocompatibility complex class I (MHC-I complex; genes HLA-A, B and C), while several components of the peptide loading complex, including TAPBP (tapasin), PDIA3 (ERp57) and CALR (calreticulin) also seemed enriched. This is consistent with a recent report suggesting that ORF8 binds to and deregulates surface expression of MHC-I molecules^26^; Fig. 2d.

### BioID proximity interactomes suggest modes of action for multiple virus proteins

In addition to providing localization information for each viral protein, individual proximity interactomes may be used to infer specific interactions that inform their function.

#### NSP7 and NSP8

In *Coronaviridae*, RNA-templated RNA synthesis is executed by two enzymes, the canonical NSP12, which employs a primer-dependent initiation mechanism^27^ and, unique to CoVs, NSP8, which is thought to provide low fidelity polymerase activity and has been suggested to work as a primase^28^. NSP7 and NSP8 associate with one another (and with NSP12), but it remains unclear how each of these individual proteins interact with the host proteome. NSP7 and NSP8 did not display particularly informative localizations by immunofluorescence or BioID compartment enrichment analysis (Extended Data Fig. 2 and Fig. 2a-b); however there is mild enrichment for a single GO term associated with the cell cycle (GO:0010564, regulation of cell cycle process, p-value 0.002 when proximity interactors for NSP7 and NSP8 are jointly analyzed). Indeed, NSP7/8 proximity interactors included several components of the MCM helicase, cohesin/condensin components, and proteins linked to the DNA damage checkpoint, histone binding, centrosome, and the anaphase-promoting complex. Interactors also included proteasome components, E1 and E3 ubiquitin system proteins, and an aminopeptidase implicated in MHC-I presentation, NPEPPS (Fig. 3a). In particular, the proximity interactors of both NSP8_Nt and NSP8_Ct were remarkably similar, suggesting that these results are not an artefact of tagging. Only a handful of the interactions reported here were previously detected; NPEPPS with NSP7, and HECTD1, HERC1, PSMD2 and UBA1 with NSP8^6,7^. Our data therefore provides new avenues of investigation for these essential virus proteins.

**Fig. 3:**
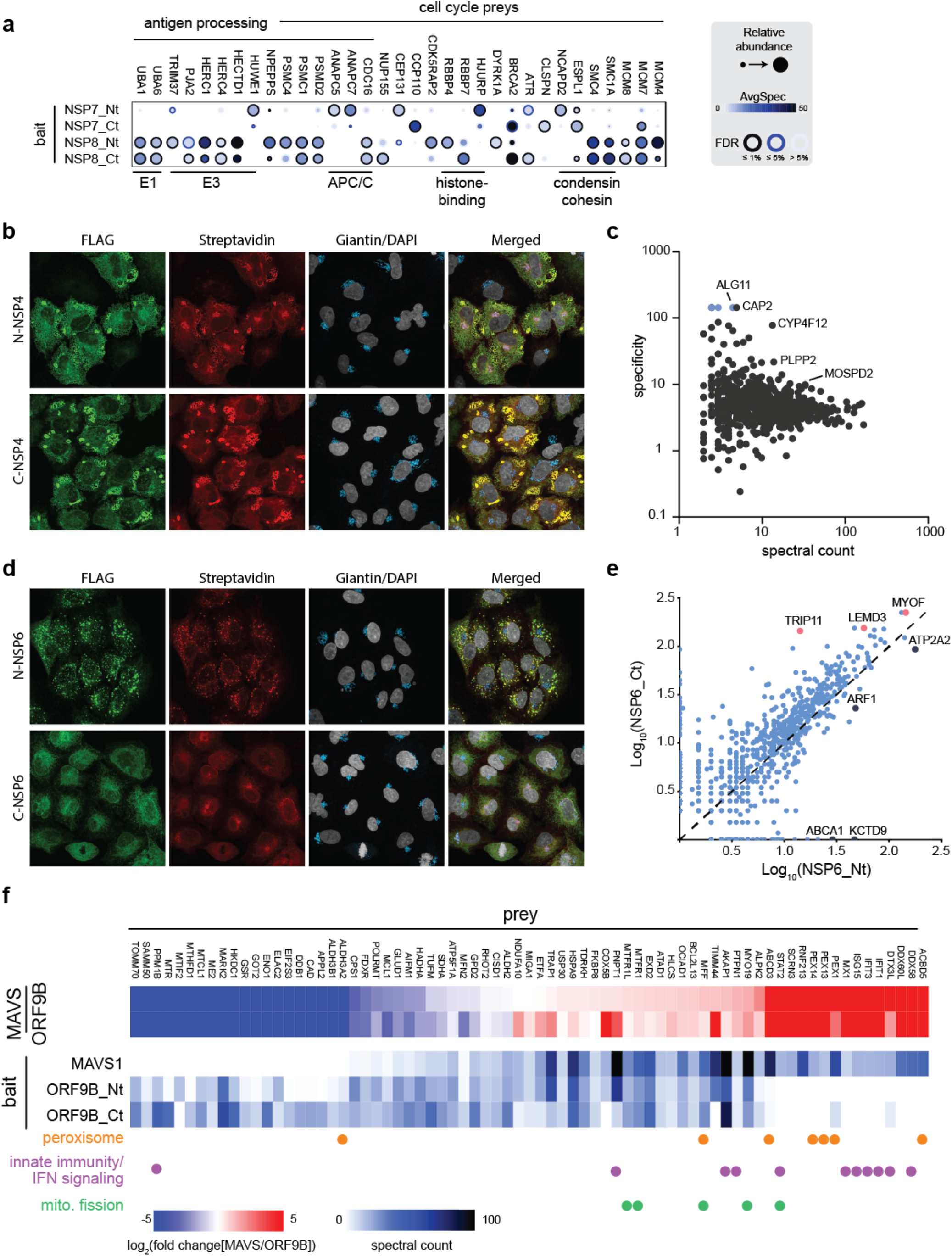
Unique localization or proximity interactomes for SARS-CoV-2 baits. **a**, Dotplot of the proximity interactors of NSP7 and NSP8 associated with the cell cycle and antigen processing (subset of 147 total preys). **b**, Phenotypes associated with the expression of NSP4, as assessed by immunofluorescence microscopy with the indicated antibodies. **c**, Specificity profile: control subtracted spectral counts (x axis) versus specificity (i.e. fold-enrichment, y axis) for preys in the NSP4_Ct BioID. The most abundant and/or specific preys are highlighted. **d**, Localization (and phenotypes) of NSP6_Nt and NSP6_Ct. **e**, Bait-bait comparison (control-subtracted spectral counts) between NSP6_Nt and NSP6_Ct. **f**, Upper heat map: fold change in spectral counts for preys detected with MAVS relative to ORF9B. (For preys with FDR of at least 0.01 and control-subtracted spectral counts of at least ten for either MAVS or ORF9B). Only preys with a consistent direction of change between ORF9B_Nt and ORF9B_Ct are shown. Fold change was capped at +/− 5 since many values are infinite (i.e. only detected with MAVS or ORF9B). Preys in the image are sorted by ORF9B_Nt and then by prey name when preys have equivalent fold changes. Lower heatmap: control-subtracted spectral counts for each indicated bait-prey pair. Prey annotated to the peroxisome, innate immunity / interferon signaling and mitochondrial fission are highlighted with colored dots.

#### NSP4 and NSP6

NSP4 and NSP6 of SARS-CoV and other coronaviruses were previously implicated in the formation of double-membrane vesicles^29,30^, a process that is critical for viral genome replication and transcription. Consistent with these observations, expression of SARS-CoV-2 NSP4, tagged at either terminus, resulted in regions with irregular membrane morphology and the formation of large vesicular structures in A549 cells (Fig. 3b). While the molecular details of this phenomenon await characterization, it is noteworthy that one of the most abundant proteins specifically enriched with NSP4 over the other viral proteins is MOSPD2 (Fig. 3c), recently identified as a scaffolding protein for ER membrane contact sites^31^. Whether overexpression of NSP4 (or viral infection) disrupts ER integrity through MOSPD2 remains to be determined. Surprisingly, despite very similar interactomes (Fig. 2b), N-terminally tagged NSP6 was localized to small cytoplasmic punctae, while C-terminally tagged NSP6 localized to the ER (Fig. 3d). Both NSP6_Ct and NSP6_Nt predominantly recover ER and Golgi proteins, suggesting that the two types of structures seen by microscopy are related. It is tempting to postulate that some of the differential interactors (Fig. 3e, Extended Data Fig. 7) may be responsible for this apparent formation of cytoplasmic punctae. Candidates could include proteins previously linked to membrane and trafficking dynamics (e.g. MYOF and ARF1), calcium or lipid signaling molecules (ATP2A2, ABCA1) or perhaps proteostasis regulators (KCTD9, USP19). Overall, the proximal interactome maps of NSP4 and NSP6 can serve as a resource to decipher the respective roles of these proteins in membrane reorganization and formation of viral replication organelles.

#### ORF9B

Consistent with its localization to mitochondria (Extended Data Fig. 2) and a proximity interactome most similar to mitochondrial proteins (Fig. 2b), ORF9B yielded a unique interaction profile enriched for mitochondrial proteins (GO CC p_adj_ 1.29e^−28^ for the C-term and 3.10e^−157^ for N-term interactomes, respectively). Tagged at either terminus, ORF9B recovered a number of outer mitochondrial membrane proteins, including TOMM70, a mitochondrial import receptor subunit shown to associate with ORF9B in three other studies^6,7,32^, and whose binding can be recapitulated *in vitro* by purified proteins^32^. ORF9B also recovers many peptides for MAVS, a protein required for innate immune defense against viruses (reviewed in ^33^) that we previously studied in the context of a BioID map of mitochondrial organization^34^. This is consistent with a recently confirmed role of ORF9B in the regulation of innate immunity^35^.

We next compared the ORF9B proximity interactome to that of MAVS: while both proteins recovered a large subset of common proximity partners (Fig. 3f), there were some notable differences. Most strikingly, proteins that were ≥2 fold more abundant in the MAVS interactome were enriched for components of the mitochondrial fission machinery, the peroxisome, and the antiviral interferon response. This observation suggests that in the absence of viral infection, ORF9B either associates with a population of MAVS excluded from peroxisomes and innate immunity regulators, or that it may interfere with the ability of MAVS to interact with these proteins. Testing these non-mutually exclusive hypotheses will require profiling of the MAVS proximal interactome following expression of ORF9B.

### Host cell RNA binding

Viruses must co-opt the host protein synthesis machinery to produce virions, and have evolved a number of elegant strategies to do so (reviewed in ^36^). While several SARS-CoV-2 bait proteins recovered ribosomal components, potentially as a consequence of their association with the ER (M, NSP2, NSP5, NSP13, NSP14 and ORF3A; Supplementary Table 1), other baits (N, NSP1, NSP3, NSP9 and NSP16) displayed a more specific recovery of cytosolic RNA-binding proteins or their interaction partners. These observations may provide important clues regarding the molecular mechanisms used by SARS-CoV-2 to effect host protein synthesis shutoff and/or usurpation of the translational apparatus.

The NSP1 interactome was uniquely enriched for eIF3G and eIF3A, two components of the eIF3 translation initiation complex, which bridges mRNA to the small ribosomal subunit (Fig. 4a). Within the evolutionarily conserved eIF3 complex, eIF3G (Tif35p in yeast) localizes near the mRNA entry channel. Our results are in agreement with with a recent report indicating that SARS-CoV-2 NSP1 effects translational inhibition by binding to and obstructing this mRNA entry channel^37^. The recovery of the largest eIF3 subunit, eIF3A, is consistent with the localization of its C-terminal domain (CTD) near this channel^38,39^. As expected from studies of SARS-CoV, NSP1 K164A and H165A mutations prevented interactions with eIF3 and ribosomal small subunits (Extended Data Fig. 8). Interestingly, however and proximity interactions with POLA1 (which were also detected by AP-MS^6^, and modelled in silico^40^) were increased. The functional significance (if any) of these and the other gained proximity interactions with the mutant NSP1 remain to be discovered.

**Fig. 4:**
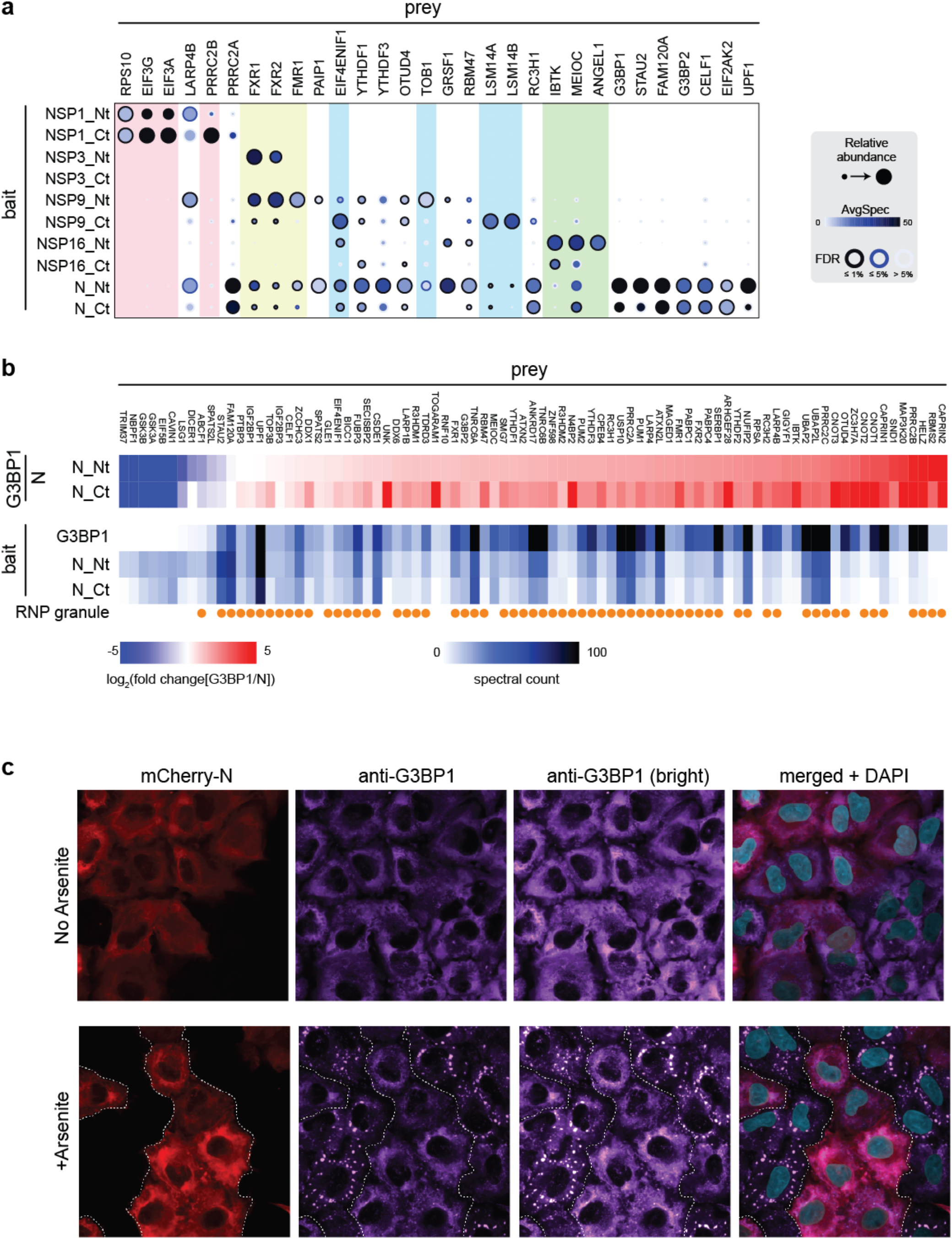
Specific RNA association and effect of N on stress granule formation. **a,** Recovery of specific cytoplasmic RNA-associated proteins (or known partners of RNA binding proteins) with the indicated baits; for N, only a subset of the RNA-related proximity partners are shown. Also see Extended Data Fig. 8. **b,** Upper heat map: fold change in spectral counts for preys detected with G3BP1 relative to N. (Preys with FDR of at least 0.01 and control-subtracted spectral counts of at least ten for either G3BP1 or N). Only preys with a consistent direction of change between N_Nt and N_Ct are shown. The fold change was capped at +/− 5 since many values are infinite (i.e. only detected with G3BP1 or N). Preys in the image are sorted by N_Nt and then by prey name when preys have equivalent fold changes. Lower heat map: control-subtracted spectral counts for each indicated bait-prey pair. Preys annotated as “Tier 1” (high-confidence) in the RNA Granule DB^49^. Other pairwise comparisons are shown in Extended Data Fig. 9). **c**. Effect of N overexpression on the formation of arsenite-induced G3BP1 stress granules. Also see Extended Data Fig. 10.

NSP3 (aka PLpro), a large (~200kDa) multi-domain polypeptide, performs proteolytic cleavage and removes ubiquitin/ISG15. While it recovered (either in its WT or proteolytically inactive form) very few partners, the RNA binding proteins FXR1 and FXR2 (related to the Fragile X mental retardation syndrome RNA binding proteins) were amongst the most abundant proximity partners of NSP3_Nt wild type and mutant proteins. Recovery of these two RNA-binding proteins and FMR1 was also detected prominently with NSP9 and N. NSP9 is a ssRNA binding protein with a unique fold^41^. In addition to the FXR/FMR proteins, NSP9 recovered several other proteins associated with RNA biology, notably LSM14A/B (implicated in the antiviral response^42^) and their direct binding partner EIF4ENIF1 (aka 4E-T^43,44^), and the deadenylase associated protein TOB1 (Fig. 4a, blue). IBTK (a protein found in immunoprecipitates of the translation initiation factor eIF4A^13,45,46^), MEOIC (a partner of the only RNA helicase-containing m6A reader, YTHDC2^47^) and ANGEL1 (a binding partner of the mRNA cap binding protein eIF4E^48^) were found in proximity to NSP16, a methyltransferase that mediates 2’-O-ribose methylation of the viral mRNA 5’-cap structure. As expected, the nucleocapsid protein, N, also established multiple proximity interactions with RNA binding proteins (a subset is shown in Fig. 4a for comparison with other baits). With the aforementioned exceptions, most of these RNA binding proteins were recovered with N more efficiently than with any of the other baits.

The N protein proximity interactome was significantly enriched for proteins localizing to the ribonucleoprotein granule GO category (GO CC P_adj_ = 0.0034). The stress granule nucleators G3BP1 and G3BP2 were amongst the most abundant N binding partners in either tag orientation (consistent with all three AP-MS studies^6–8^; other previously reported interactions include those with UPF1^6^ and CAVIN1/2^7^). Using classic BioID (BirA* in HEK293 cells), we previously reported a proximity interactome of cytoplasmic RNA granules and bodies^13^. One of these structures, the P-body, contains components of the decapping machinery (including DCP1A) and deadenylase complexes, and is present as visible structures by microscopy in untreated cells. Another of these structures, the stress granule, is marked by proteins such as G3BP1, and forms as a microscopically distinguishable structure only after cells are subjected to stress (e.g. exposure to sodium arsenite). While DCP1A and G3BP1 provide relatively clean definitions of P-bodies and stress granules by microscopy, several proteins associate with both structures by microscopy and BioID (reviewed in ^49^). These include LSM14A, a main component of P-bodies, but which is also found in stress granules. GRSF1, the first protein identified as a mitochondrial RNA granule component, also establishes proximity interactions with cytoplasmic granules.

Interestingly, we noted obvious differences in the recovery of proximity partners for N versus G3BP1 (Fig. 4b), LSM14A, GRSF1 and DCP1A (Extended Data Fig. 9). In particular, CAPRIN1 and UBAP2L (and its paralog UBAP2), which bind to G3BP1 and are necessary for nucleating stress granules^13,50^, were proportionally depleted in the N BioID in comparison to that of G3BP1 BioID (Fig. 4b), suggesting that N overexpression could oppose G3BP1-mediated stress granule formation. To test this hypothesis, we investigated the impact of the expression of N (tagged with mCherry) on the formation of stress granule foci (detected by endogenous G3BP1) upon treatment with sodium arsenite. After 30 min of arsenite treatment, G3BP1 forms punctae in cells not expressing N protein. However, expression of N drastically decreased the formation of these structures in response to arsenite (Fig 4C and Extended Data Fig. 10). Interestingly, this phenomenon was rescued upon co-expression of GFP-tagged G3BP1, suggesting that N could titer functional G3BP1 away from its stress granule co-nucleators (note that in the context of G3BP1 overexpression, N localized to stress granules; Extended Data Fig. 10). Taken together, these results suggest that a new function for N may be to prevent the formation of host cell stress granules, likely by competing with the association of proteins such as UBAP2, UBAP2L and CAPRIN1 that promote stress granule formation.

## Discussion

Here we report the proximal proteome of SARS-CoV-2 proteins, profiled as both N- and C-terminal fusions with the fast acting miniTurbo enzyme, in A549 cells, a widely used model system in coronavirus biology^50–52^. We present the dataset as a resource accompanied by a user-friendly website (covid19interactome.org), and apply immunofluorescence microscopy, term enrichment and bait similarity profiling to assign each bait to a specific subcellular location. The resulting dataset identifies a number of new and confirmatory proximity interactions, and can be mined to shed light on the viral life cycle and to identify virus-host interactions that could represent important therapeutic targets.

A major challenge in the interpretation of proximity-dependent biotinylation is the complexity of the data that it generates^9^. While some of the bait proteins (NSP1, for example; Extended Data Fig. 8) yielded a short list of proximity partners that immediately suggested mechanistic insights, this is not always the case. In particular, baits localized to membranes recovered hundreds of proximity partners. While it may be tempting to report that the most abundant proximity partners detected in these analyses are more closely associated with the bait, this may not always be true. The extent of biotin labeling and recovery of each prey protein is dependent on a variety of factors, including its size, amino acid composition, abundance and propensity for biotinylation (i.e. the number of unmodified lysine residues on a given prey protein that are available for labeling, in the context of the specific cellular compartment). It is also important to note that a given bait protein may have multiple *bona fide* intracellular locations. In this case, some prey proteins with relatively low peptide counts can represent critically important interactors that provide extremely useful information on bait function and cellular context. To assist in data interpretation, we have leveraged our human cell map resource (humancellmap.org) to select host baits that profile different subcellular localizations. This data was very helpful for more clearly characterizing local virus bait environments, and allowed us to better understand more complex virus protein interactomes (as shown when we e.g. compared the ORF9B and MAVS interactomes, and compared the N interactome with those of G3BP1 and other RNA granule components). In particular, comparing virus and host marker protein interactomes allowed us to: (i) better refine modes of targeting to a particular subcellular structure (e.g. likely through TOMM70 for ORF9B targeting to the mitochondria) and; (ii) provided testable hypotheses regarding the functional consequences of these virus-host associations.

Applying this conceptual framework to SARS-CoV-2 protein N and host protein G3BP1, and in particular noting the relative depletion of proteins critical to the formation of stress granules (UBAP2L, its paralog UBAP2, and CAPRIN1 in the N BioID interactome), we postulated that N may bind to G3BP1 to interfere with the association of UBAP2L/2 and/or CAPRIN1, and potentially prevent stress granule formation. This hypothesis was further bolstered in our microscopy experiments. While future work will be required to determine whether this phenomenon results from direct competition between UBAP2L (and/or CAPRIN1) and N for a common binding site on G3BP1, this observation may be highly relevant to understanding the SARS-CoV-2 life cycle.

An important component of the host response to viral infection is the activation of the kinase PKR (gene name EIF2AK2), which phosphorylates the translation initiation factor eIF2alpha at Ser51, resulting in translational arrest and the formation of stress granules^53^. Multiple viruses have evolved strategies to interfere with this process, either through preventing PKR activation and eIF2alpha phosphorylation, or (as is being increasingly recognized) by modulating stress granule dynamics by other means. For example, the poliovirus 3C protease was shown to cleave G3BP1^53^. How the recent characterization of a phase-separation phenomenon associated with N and viral RNA^54–57^ relates to our findings should be addressed in the context of viral infection, or at least viral RNA transfection.

Several recent AP-MS studies have identified biochemically-stable interactions for SARS-CoV-2 proteins. As with previous comparative studies^11,12,15^, the overlap between the AP-MS and BioID datasets varies depending on the bait. This is not surprising, as, in addition to major differences in methodologies, BioID is a proximity-dependent detection method, and thus simply captures proteins in the same local environment as the bait. AP-MS conducted under standard, gentle lysis conditions is not optimal for the capture of labile protein-protein interactions, or proteins associated with membranes. Post-lysis artefacts, or in some cases associations that result from the extreme overexpression of transient transfection, can also explain some of the interactions captured by published AP-MS studies that are not recapitulated here. On the other hand, false negatives in BioID experiments can relate to the requirement for solvent-exposed lysines, and for sufficient engagement time to enable biotinylation. Lastly, the high stringency employed in our BioID experiments is expected to dissociate most protein-protein interactions, so that only those proteins marked by biotin will be identified: this results in a selective loss of stable but non-biotinylated components of protein complexes in our studies. Given these caveats, the detection of ~21% of the previously-reported AP-MS interactions (428) is not unexpected. Importantly, given the orthogonality of the approaches, hits detected with high confidence with both approaches are much more likely to represent bona fide interactions, and some illustrative examples are included in this manuscript.

There are limitations to the current study. First, as mentioned above, any epitope tagging may influence the behavior of the tagged protein, possibly affecting its localization and/or interactions. Here, we have attempted to mitigate this variable by separately tagging the N- and C-termini. This, however, still left some unanswered questions, for instance regarding the localization and proximity interactors of ORF14 (also called ORF9c in other studies). We have analyzed proximity interactomes for viral proteins expressed individually in uninfected cells. This is useful to illuminate individual connections of each viral protein with the host proteome, but it does not reveal new connections that could be formed by the association of viral proteins with one another. For viral proteins that dimerize (e.g. NSP7-NSP8), the next step could be to analyze the proximal proteome of the dimer rather than the monomer, a task that can be facilitated by using a Protein Complementation Assay strategy for BioID (as in ^58^). Ultimately, though, it will be critical to profile the proximal interactome of each viral protein in the context of a SARS-CoV-2 infection; this will also be particularly important for understanding the formation of viral factories.

While the majority of hospitalized COVID-19 patients present with compromised airway function, a wide variety of effects on other organs have been noted. The primary SARS-CoV-2 receptor, angiotensin converting enzyme 2 (ACE2^59^), is expressed in a wide variety of cell types^60^, including cardiomyocytes, cardiac pericytes, endothelial cells, fibroblasts, hepatocytes and multiple cell types in the kidney^61^, and kidney impairment^62,63^, cardiac injury^64^ and liver dysfunction^65,66^ are commonly observed in COVID-19 critical care patients. While the virus-related cytokine storm has been linked to organ damage^67^, a recent report also indicated that SARS-CoV-2 virus RNA was found in autopsied lung, pharynx, heart, liver, kidney, and brain^68^. Our virus-host proximity map of a lung-derived cell line provides a comprehensive proximity interaction map for this model system, but the proteome of different cell types can vary significantly, and virus tropism and replication efficiency will likely vary dramatically across cells derived from different tissues. It will thus be of critical importance to characterize virus-host proximity interactomes in multiple human tissues/cell types. Our lentiviral delivery system for inducible expression of viral proteins will greatly facilitate cell type cross-comparisons, as well as comparisons of viral proteins from related coronaviruses, which could shed light on the unique properties of SARS-CoV-2. Lastly, use of the fast acting miniTurbo enzyme (which allows for a labeling duration of ≤15 minutes as compared to ~6 or more hours for BirA*) enables us to both generate more temporally resolved profiles, and will allow us in future to track changes in virus-host proximity interactions following drug treatment, or to follow changes in associations throughout the viral life cycle.

## Acknowledgements

**People**: We thank members of our laboratories for helpful discussions, Guomin Liu for assistance with mass spectrometric analysis and Daniel Schramek for providing cell lines.

## Author Contributions

PST and HA generated all the reagents for BioID from Gateway entry constructs generated by DYK, JJK, PC, CD, YJ, FPR and BR.

ZYL performed BioID purification and ran the mass spectrometer with help from BL and CJW PST, HA, AA, RS and JSG performed experiments

PST, JDRK and ACG analyzed data

JDRK created the website

ACG, FPR, YJ and BR acquired funds for this project and supervised team members

ACG, BR, JDRK and PST wrote the manuscript, with input from other authors

## Materials and Methods

### Cloning

The pSTV2-3xFLAG-miniTurbo vector was generated by modifying the previously described pSTV2-FLAG-BirA* lentiviral transfer vector^18^. Briefly, the human codon optimized sequence for miniTurbo^17^ was synthesized together with 3 copies of the FLAG epitope and a Glycine-Serine flexible linker and inserted into the pSTV2-FLAG-BirA* backbone.

Open Reading Frames for the SARS-CoV-2 predicted proteins reported elsewhere^19^ were subcloned by Gateway cloning into the pSTV2-miniTurbo N- and C-termini containing vectors. Point mutants in NSP3 (C587A) and NSP5 (C145A) that abrogate protease activity and in NSP1 (K164A and H165A) as in ^22^ were constructed by site-directed mutagenesis.

Entry vectors for host cell proteins previously used as proximity-dependent biotinylation (BioID) subcellular markers were transferred into pSTV2-miniTurbo N- or C-termini vectors (the orientation was kept consistent with our previous study^20^). MAVS entry vector was subcloned with a N-terminal tag as in ^34^. The signal sequence from IgKV4-1 (MVLQTQVFISLLLWISGAYG) followed by an HA-tag (YPYDVPDY) and residues 617-675 from CD44 that contains a portion of the extracellular domain, and the transmembrane domain (residues 643-675) without a stop-codon were amplified using standard AttB1/2 gateway primers to generate an entry vector. All entry and destination vectors were validated by restriction digests and/or sequencing and all resulting expression vectors were confirmed by restriction digestion and functionally validated by transient transfection and immunostaining to confirm bait expression and miniTurbo functionality.

Entry clones for G3BP1 and the SARS-CoV-2 nucleocapsid protein (N) were cloned into pSTV6H Gateway destination vectors containing an in-frame N-terminal EGFP or mCherry tag. pSTV6H is similar to the previously described pSTV6 vector^18^, however, the PGK promoter was replaced with a minimal EF1a-promoter.

### Arsenite Treatment and Stress Granule Formation

A549 cells expressing SARS-CoV-2 Nucleocapsid protein tagged with mCherry with and without GFP-tagged G3BP1 were plated on coverslips and expression induced using 1μg/ml doxycycline for 24 hours. Cells were treated with 0.5 mM sodium arsenite (Fluka; 35000-1L-R) for 30 minutes to induce stress granule formation prior to fixing and staining (see Immunofluorescence methods).

### Cells, transfection and infection

HEK293TN cells (a gift from Daniel Schramek) were maintained in DMEM high glucose with 10% heat-inactivated FBS and 1% Penicillin/Streptomycin and used for virus production. A549 cells (a gift from Daniel Schramek) were maintained in DMEM high glucose with 10% heat-inactivated FBS and 1% Penicillin/Streptomycin.

Lentivirus production was performed in HEK293TN cells using jetPRIME reagent as per manufacturer’s recommendations (Polyplus-transfection SA, Illkirch-Graffenstaden, France, Cat# 114 – 01) as previously^18^.

For all BioID experiments performed in A549 cells, 2 million cells were seeded in 15 cm dishes and infected with rtTA lentivirus and ORF containing pSTV2 lentivirus, resulting in 80-90% transduction efficiency. The following day, cells were split into two 15 cm plates with an aliquot of the cell suspension plated onto coverslips for immunofluorescence. The next day, the transgene was induced using 1 μg/ml doxycycline for 24 hours (in biotin-depleted media to minimize spurious proximal labeling). Cells were then treated with 50 μM Biotin for 15 minutes prior to fixing for immunofluorescence or lysis for BioID and western blot. Biological duplicates were prepared for all experiments (alongside negative controls defined below).

For the baits that posed a challenge in lentiviral production (NSP1-wt and NSP3), A549 cells were transduced with rtTA virus as described above. They were then seeded in 15 cm plates and transfected with the 4 μg of the respective miniTurbo-tagged ORF in the pSTV2 transfer vector using jetPrime reagent to boost the expression levels prior to biotinylation and harvesting as above.

### Immunofluorescence

Cells were plated on coverslips and induced with doxycycline and biotin as described above. After 24 hours, cells were washed once with PBS and fixed in 4% paraformaldehyde in PBS for 10 minutes. Subsequently, cells were washed, permeabilized with 0.25% NP-40 in PBS, and blocked in 4% skim milk in PBS. Cells were stained for bait proteins using mouse anti-FLAG antibody (Monoclonal anti-FLAG M2 antibody, Sigma-Aldrich, Cat# F3165; used at 1:2500), Rabbit-anti-Giantin antibody (Abcam, Cat# AB24586, 1:1000) and anti-G3BP1 antibody (mouse polyclonal, BD Transduction Labs, Cat# 611126; used at 1:500), in blocking buffer. Secondary detection of bait proteins was performed in 2.5% bovine serum albumin (BSA) in PBS using goat anti-mouse Alexa FluorTM 488 (Molecular Probes, ThermoFisher Scientific, A11001, used at 1:1000) and Alexa FluorTM 594 streptavidin conjugate (Molecular Probes, ThermoFisher Scientific, Cat# S11227, used at 1:2500) was used to localize the sites of in vivo biotinylation. DAPI (Sigma Aldrich, 20 mg/ml, used at 1:10,000) was used as a nuclear counterstain. Slides were mounted in ProLong Gold AntiFade (Molecular Probes, ThermoFisher Scientific, Cat# P36930) and imaged on a Nikon A1R+HD confocal scanner attached to an Eclipse Ti2-E inverted microscope. Confocal scanning was performed with a resonant scanner and 60x/1.4 CFI Plan Apo lambda oil objective. NIS-Elements software was used for image acquisition and processing with Denoise.AI and 3D deconvolution. Images were processed using Volocity software V6.2 (PerkinElmer, Waltham, MA).

### Immunoblotting

Cells were lysed as described for BioID (see below) and boiled in Laemmli SDS-PAGE sample buffer. Proteins were resolved on 4–15% Criterion™ TGX™ Precast gels (Bio-Rad Laboratories Inc., Cat# 5671085) and transferred to nitrocellulose membranes (GE Healthcare Life Science, Uppsala, Sweden, Cat# 10600001) for immunoblotting. Following Ponceau S staining, membranes were blocked in 4% skim milk in TBS with 0.1% Tween-20 (TBST). Bait proteins were probed using mouse anti-FLAG antibody (Monoclonal anti-FLAG M2 antibody, Sigma-Aldrich, F3165; used at 1:2500) in blocking buffer, washed in TBST and detected with Sheep anti-Mouse IgG-Horseradish peroxidase (HRP; GE Healthcare Life Science, Cat#NA931; used at 1:5000). Similarly, biotinylated proteins were probed using HRP-conjugated Streptavidin (GE Healthcare Life Science, Cat# RPN1231vs; used at 1:2500) in 2.5% bovine serum albumin (BSA) blocking buffer. Membranes were developed using LumiGLO chemiluminescent reagent (Cell Signaling Technology, Danvers, MA, Cat# 7003S).

### BioID

Cells at 75% confluence in 15 cm plates were induced with 1 μg/ml doxycycline (Dox) for 24 hours at which point biotin was added to the media at a final concentration of 50 μM for 15 minutes. Cells were washed twice with cold PBS on ice and lysed on plate using 1 ml of modified RIPA (modRIPA) buffer [50 mM Tris-HCl, pH 7.4, 150 mM NaCl, 1 mM EDTA, 1 mM MgCl2, 1% NP40, 0.1% SDS, 0.4% sodium deoxycholate] containing 1 mM PMSF, 1x Protease Inhibitor mixture (Sigma-Aldrich, Cat# P8340), 25 U of TurboNuclease (BioVision Inc., Milpitas, CA, Cat# 9207) and 10 μg of RNase A (Bio Basic, Markham, ON, Canada, Cat# RB0473) and incubated on ice for 5 minutes with occasional agitation of the plate. Subsequently, 10% SDS was added to the plate to raise the SDS concentration to 1% SDS at which point the lysate was scraped, transferred to an 1.5 ml tube, flash frozen on dry ice and frozen at −80 °C for downstream processing.

For affinity purification, lysates were thawed at 4 °C, and sonicated for 15 seconds (5 seconds on, 3 seconds off for three cycles) at 30% amplitude on a Q500 Sonicator with 1/8” Microtip (QSonica, Newtown, Connecticut, Cat# 4422). To reduce and alkylate the samples, DTT was added to each tube to a final concentration of 5 mM and incubated at 60 °C for 30 minutes. Iodoacetamide was then added to a final concentration of 10 mM and the samples were incubated in the dark at room temperature for 20 minutes. Samples were centrifuged at 15,000 × g for 15 minutes and the supernatant was used for biotinylated protein capture using 15 μl of pre-washed Streptavidin agarose beads (GE Healthcare Life Science, Cat# 17511301). After 6 hours, the beads were transferred to a new tube, washed once with SDS-Wash buffer (25 mM Tris-HCl, pH 7.4, 2% SDS), once with RIPA (50 mM Tris-HCl, pH 7.4, 150 mM NaCl, 1 mM EDTA, 1% NP40, 0.1% SDS, 0.4% sodium deoxycholate), once with TNNE buffer (25 mM Tris-HCl, pH 7.4, 150 mM NaCl, 1 mM EDTA, 0.1% NP40), and two times with 50 mM ammonium bicarbonate (ABC buffer), pH 8.0, and finally with 50 mM ABC buffer supplemented with 0.0025% ProteaseMax (Promega, Cat# V2071) to facilitate sedimentation of agarose resin. On-bead digestion was performed overnight at 37 °C with 0.75 μg of trypsin (Sigma Aldrich, Cat# 6567) in 60 μl of ABC buffer, followed by further digestion with an additional 0.25 μg of trypsin for 3 hours. Peptide supernatants were collected into a new tube, beads were washed twice with HPLC grade water, pooled with the peptide supernatant, acidified by adding 0.1 volume of 50% formic acid, and subsequently dried using vacuum centrifugation.

### Mass Spectrometry

The LC-MS/MS setup consisted of a TripleTOF 6600 (SCIEX, Framingham, MA, Canada) equipped with a nanoelectrospray ion source connected in-line to a 425 Nano-HPLC system (Eksigent Technologies, Dublin, CA). The fused silica column (15 cm × ID 100 μm, OD 360 μm) had an integrated emitter tip prepared in-house using a laser puller (Sutter Instrument Co., Novato, CA). The column was packed with ∼15 cm of C18 resin (Reprosil-Pur, 3 μm, Dr. Maisch HPLC GmbH, Germany). Lyophilized samples were resuspended in 5% formic acid and 1/4th of sample was used per injection for DDA and 1/4th for DIA. IRT calibration mixture (Biognosys (Zurich, Switzerland)) was added to the mixture for calibration of retention times. Samples were loaded onto the column using the autosampler, and the LC delivered the organic phase gradient at 400 nl/min over 90 min (2–35% acetonitrile with 0.1% formic acid). The MS instrument was operated in data-dependent acquisition mode with 1 MS scan (250 ms; mass range 400–1250 m/z) followed by up to 20 MS/MS scans (50 ms each). Only candidate ions between two and five charge states were considered, and ions were dynamically excluded for 10 s with a 50 mDa window. The isolation width was 0.7 m/z, and minimum threshold was set to 200. Between sample injections, 2 blank samples were injected (5% formic acid), each with 3 rapid gradient cycles at 1500 nl/min over 30 min. Before another sample was injected, system performance was verified with a 30 min BSA quality control run and a 30 min BSA mass calibration run.

### Peptide and Protein Identification

Raw files (.WIFF and .WIFF.SCAN) were converted to an MGF format and to an mzML format using ProteoWizard (v3.0.4468) and the AB SCIEX MS Data Converter (V1.3 beta), as implemented within ProHits^69^. For human samples, the database used for searches consisted of the human and adenovirus sequences in the RefSeq protein database (version 57). The database was supplemented with SARS-CoV-2 protein sequences, “common contaminants” from the Max Planck Institute (http://141.61.102.106:8080/share.cgi?ssid=0f2gfuB) and the Global Proteome Machine (GPM; http://www.thegpm.org/crap/index.html), and with commonly used epitope tags. The search databases consisted of forward and reverse sequences (labeled “gi 9999” or “DECOY”); in total, 72,226 entries were searched for the human database. Spectra were analyzed separately using Mascot (2.3.02; Matrix Science) and Comet^70^ [2012.01 rev.3] with trypsin specificity and up to two missed cleavages; deamidation (Asn or Gln) and oxidation (Met) were selected as variable modifications and carbamidomethylation of cysteine residues was selected as a fixed modification. The fragment mass tolerance was 0.15 Da, and the mass window for the precursor was ±40 ppm with charges of 2+ to 4+ (both monoisotopic mass). The resulting Comet and Mascot results were individually processed by PeptideProphet^71^ and combined into a final iProphet^72^ output using the Trans-Proteomic Pipeline (TPP; Linux version, v0.0 Development trunk rev 0, Build 201303061711). TPP options were as follows: general options were -p0.05 -x20 -PPM -dDECOY, iProphet options were -pPRIME, and PeptideProphet options were -pPAEd. All proteins with a minimal iProphet probability of 0.95 were used for analysis.

### Identification of High-confidence Proximity Partners

SAINTexpress^23^ (version 3.6.1) was used to calculate the probability that identified proteins were enriched above background contaminants. SAINTexpress uses a semi-supervised spectral counting model that compares the detection of putative proximal interactors in a BioID profile of a given bait against a series of negative control runs. For analysis with SAINT, only proteins with an iProphet protein probability of >0.95 were considered, and a minimum of two unique peptides was required. For each cell type, bait proteins were profiled using independent biological duplicates and analyzed alongside six independent negative control runs from the same cell type. Negative control runs consisted of streptavidin purifications from cells expressing miniTurbo-EGFP only (this models promiscuous biotinylation), miniTurbo-NES (nuclear exclusion signal) or without miniTurbo transgene expression (this models endogenous biotinylation). For running SAINTexpress, the 12 independent negative controls were compressed to three, meaning that for each prey, its three highest counts across the 12 controls were selected for stringent evaluation with SAINTexpress (as detailed previously^73^). SAINTexpress scores were averaged across both biological replicate purifications of the baits, and these averaged values were used to calculate a Bayesian False Discovery Rate (FDR); proximity interactions detected with a calculated FDR of 1% or less were deemed of high confidence.

### SAINT processing

The SAINT file was filtered to remove contaminants, tags and viral bait genes detected as preys. Genes that are now obsolete since the Refseq version was released were also removed, as were fusion proteins, their composite genes, and protein isoforms whose peptides were not reliably assigned between replicates and/or baits. The SAINT file was supplemented with columns for the control subtracted spectral count, the control-subtracted and length adjusted spectral count (CLSC) and prey specificity. Control subtracted spectral counts were calculated as the average spectral count in controls subtracted from the average spectral count (AvgSpec), which could be no less than zero. The CLSC was calculated by taking the control subtracted spectral count for a prey and multiplying by the median prey sequence length (within each bait), divided by the sequence length of the prey in question. The specificity of a prey with a bait was calculated by dividing the control subtracted spectral count by the average control subtracted spectral count for that prey across all other bait genes. When calculating the specificity of a prey for bait **X**, other baits from the same gene as bait **X** were ignored, and other baits were consolidated by gene using the highest spectral count.

### Enrichments

Gene Ontology (GO) enrichments were performed using g:Profiler’s Python client^74^ (database version e100_eg47_p14), with the g:SCS multiple testing correction method, a significance threshold of 0.01, our defined A549 custom background set and with no electronic annotations. The A549 background set included all proteins defined in the SAINT report or those found in A549 cells in ProteomicsDB^75^ with an MS1 intensity > 0 (Supplementary Table 5).

Protein domains and regions were retrieved from Pfam^76^ and enrichments were calculated for each bait using Fisher’s exact test using the A549 background. The FDR was controlled by using the Benjamini–Hochberg procedure for an FDR of 1%. Domains had to be found in at least three preys to be considered as enriched. Allowable regions included coiled_coil, sig_p and transmembrane.

### Previously reported interactions

Known interactions for SARS-CoV-2, SARS-CoV and MERS were downloaded from BioGRID^77^ (version 3.5.188).

### Data Visualization

Dot plots and heat maps were generated using ProHits-viz^78^ (prohits-viz.lunenfeld.ca). SAINT FDR is represented as the edge color intensity. Quantitation is encoded using the color gradient representing control-subtracted spectral counts (capped at 50), with relative spectral counts across baits represented by node size.

### Network view

The network was generated using Cytoscape^79^ version 3.8.0. At most the top 25 preys for each bait were used based on the control-subtracted and length normalized spectral count (CLSC) after merging the N- and C-terminal versions of the baits. If a prey was detected for both the N- and C-terminal versions of a bait, the record with the better FDR was used when merging, or the better spectral count if the FDR was equivalent. The network was laid out using the Prefuse Force Directed Layout algorithm with the yFiles layout used to remove node overlaps.

### Experimental Design and Statistical Rationale

For each BioID experiment, biological duplicates were employed (each replicate generated through independent infections and harvests). Statistical scoring was performed against six negative controls compressed to two virtual controls using Significance Analysis of INTeractome (SAINT; SAINTexpress 3.6.1 was employed) as described above (“Identification of High-Confidence Proximity Partners”). Control samples (“No-miniTurbo”, “miniTurbo-EGFP”, “miniTurbo-EGFP-NES”) are described above. The average SAINTexpress score was used to determine the Bayesian FDR, which therefore requires a high-confidence interaction to be detected across both biological replicates.

---

**Extended Data Fig. 1.**
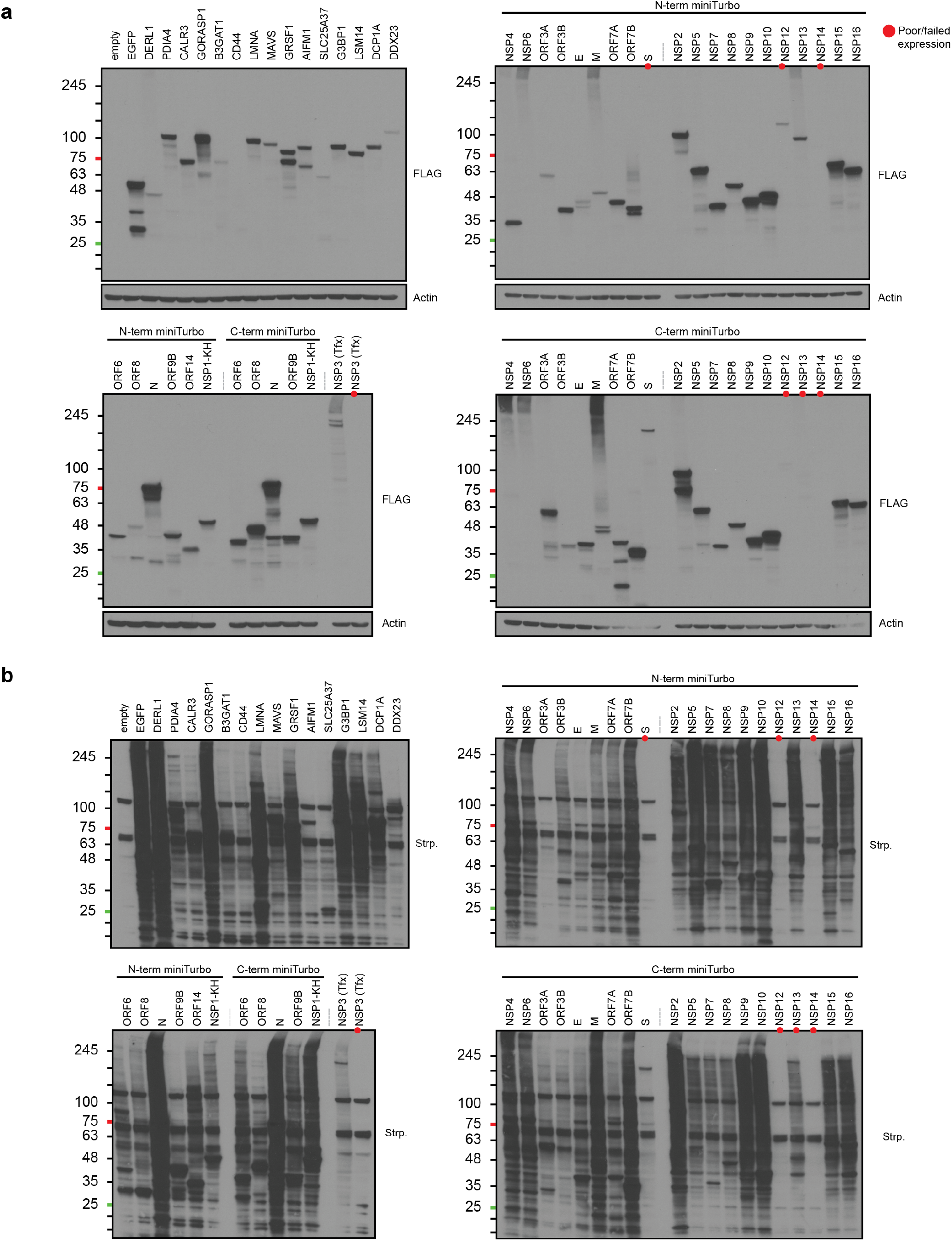
Expression of the bait proteins in A549 cells. **a-b,** Protein expression in the indicated transduced cells was induced with doxycycline (24 hours). Biotin (50 μM) was added to the culture media (15 minutes) and cells were harvested as detailed in methods. Proteins were resolved on SDS-PAGE and blotted with anti-FLAG. Beta-actin was used as loading control (**a**). Membranes were quenched using NaN3 and reprobed for biotinylated proteins using HRP-conjugated streptavidin (**b**).

**Extended Data Fig. 2a,b:**
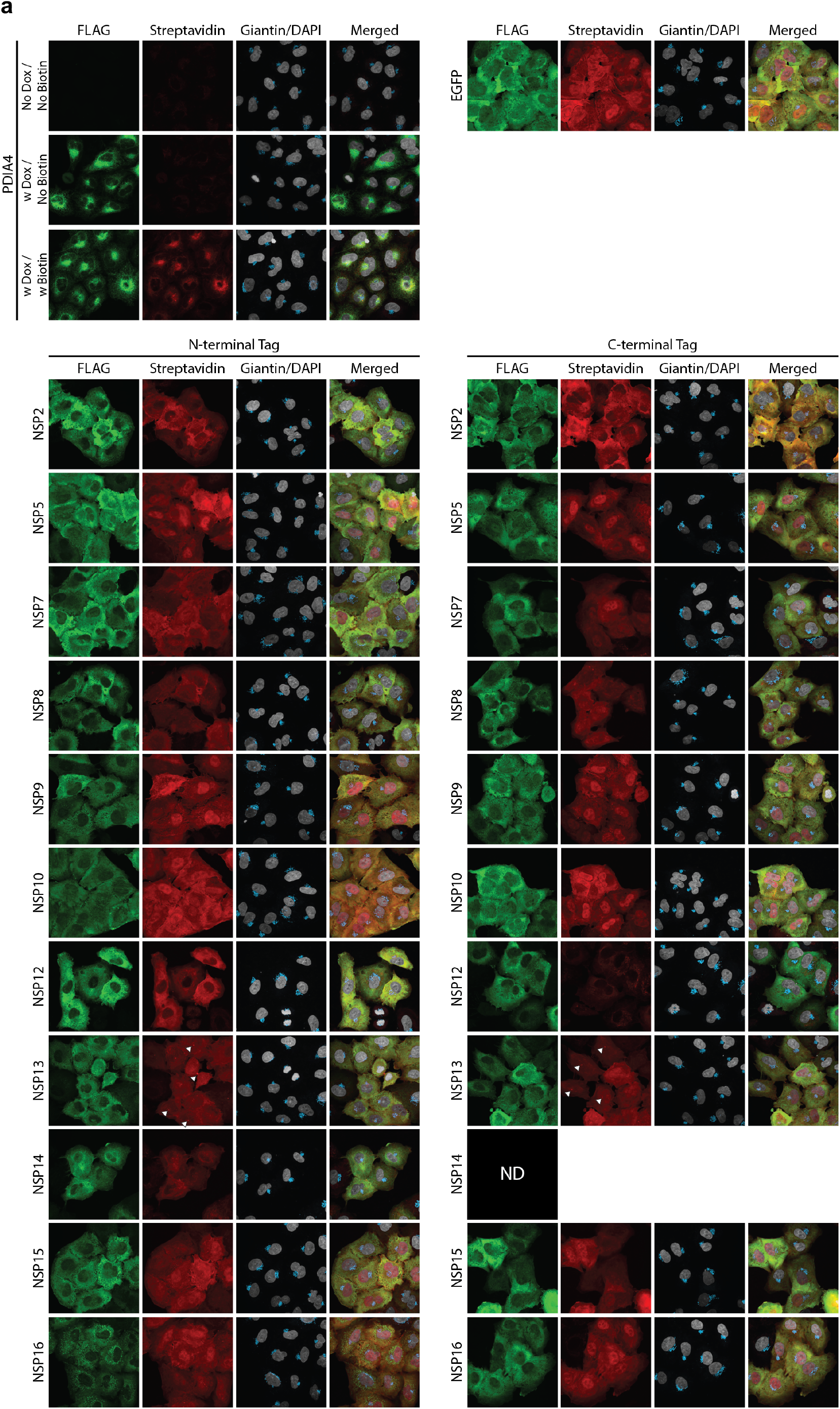
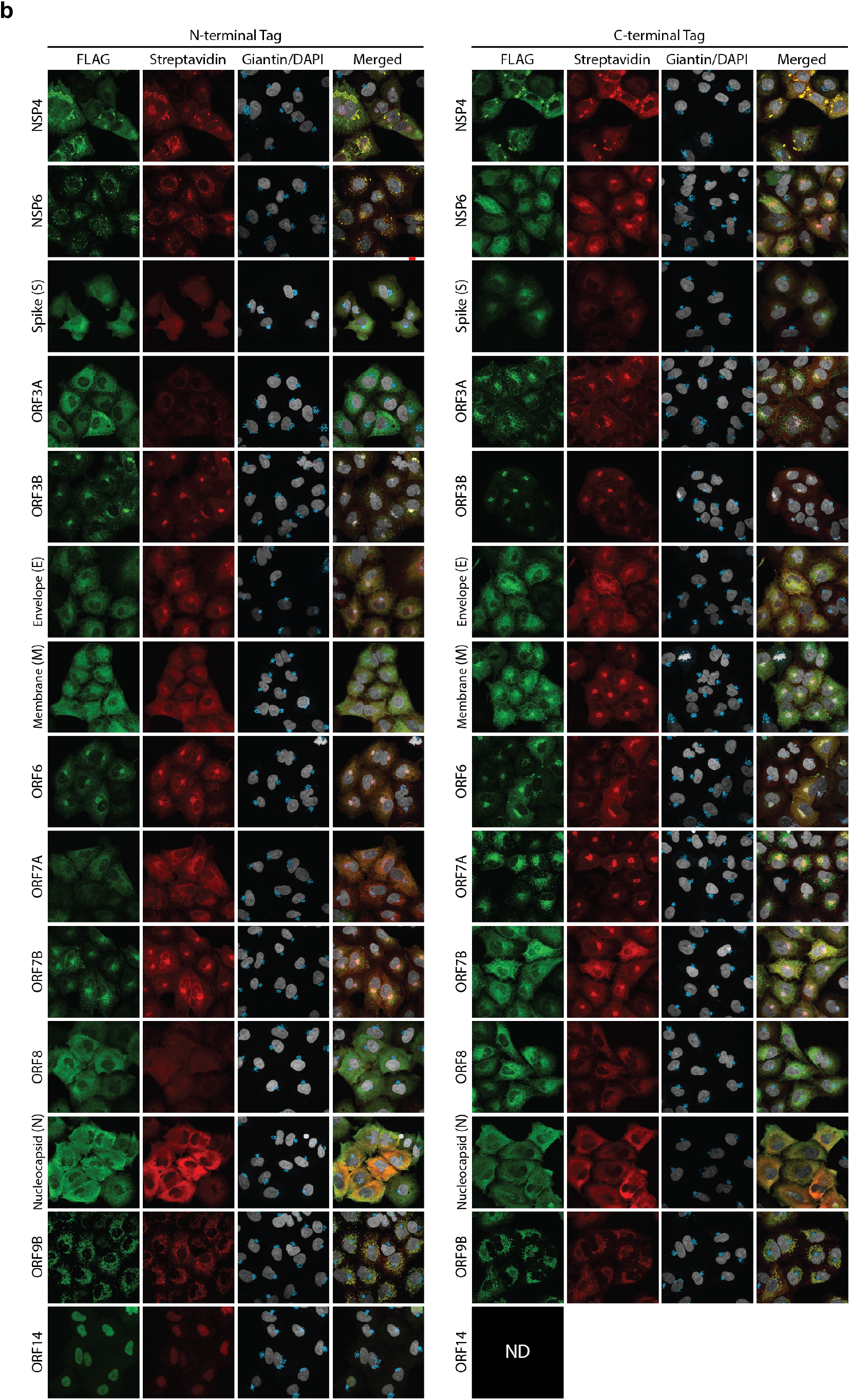
Bait protein localization and proximal labeling. A549 cells were transduced with lentivirus to deliver the specified bait protein. Bait expression was induced as above. Cells were fixed and stained for bait protein using anti-FLAG antibody and for biotinylated proteins using Alexa-594 conjugated streptavidin. The Golgi and nucleus were counterstained using anti-Giantin antibody and DAPI, respectively. Staining controls (for host bait PDIA4) included omission of doxycycline or biotin. Due to variability in expression levels between different baits, exposure settings and image brightness were adjusted to enable visualization of all baits shown, and is not intended to be quantitative. Images for NSP1, NSP3 NSP14_Ct and ORF14_Ct were not acquired (ND) due to low expression levels.

**Extended data Fig. 3:**
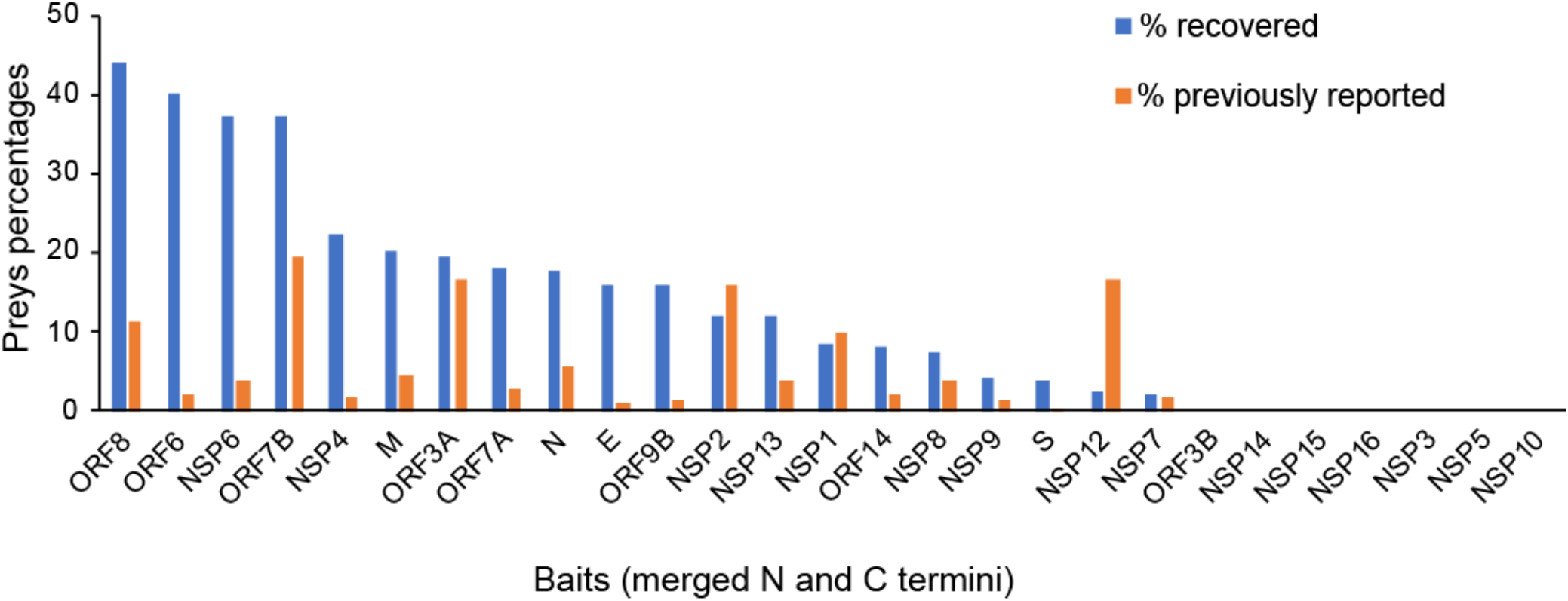
Overlap with previously reported interactions by bait (supports Fig. 1e). Percentage of previously reported interactions that were recovered by each bait in BioID, and the percentage of BioID proximity interactions that have been previously reported (See Supplementary Table 3).

**Extended Data Fig. 4.**
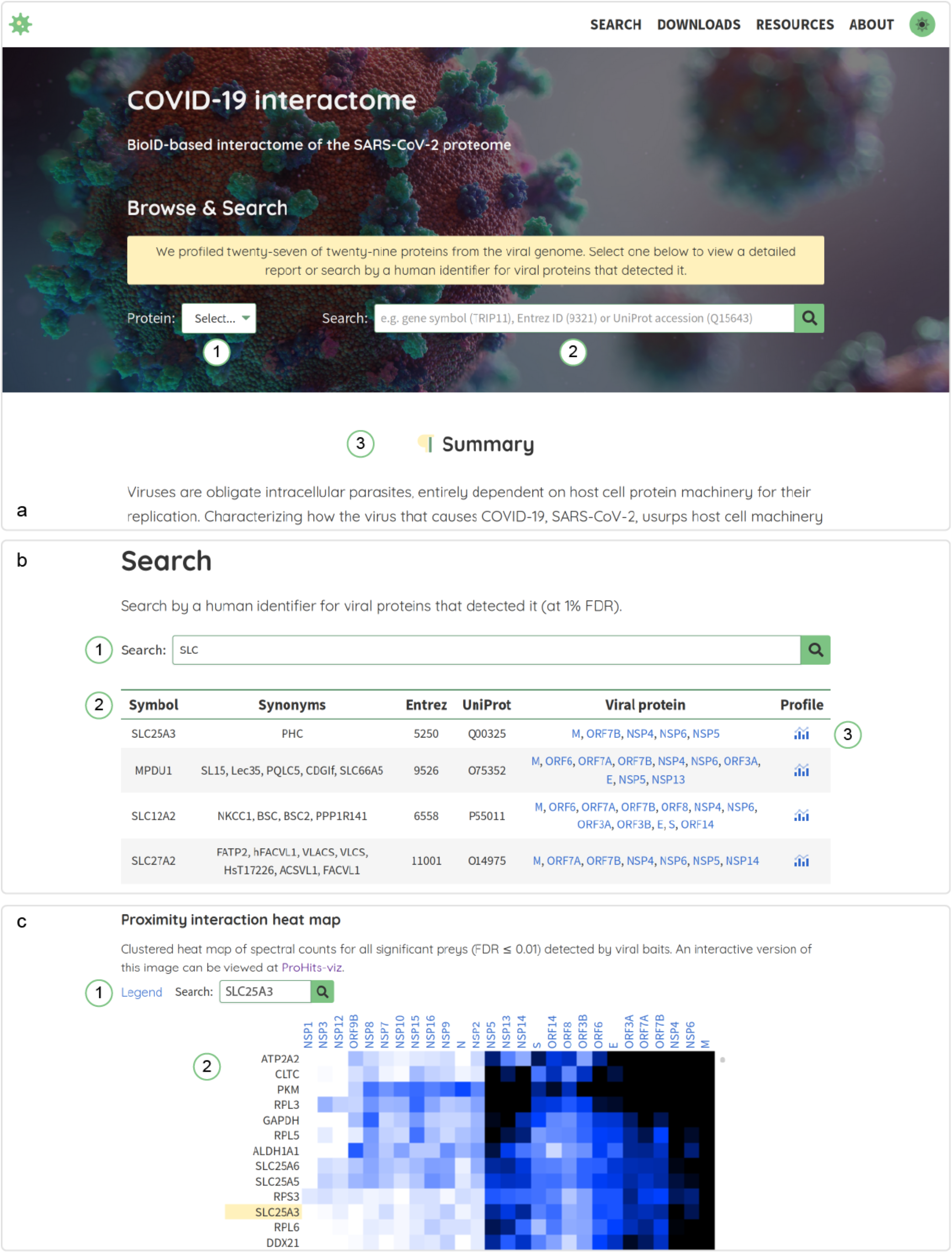
SARS-CoV-2 website. All data reported in this manuscript are available for browsing and downloading at covid19interacome.org. **a**, From the home page users can select a viral bait and view its report from (1) or search by a human gene identifier for viral baits that detected it (2). We also provide an overview of the project and its findings beneath the search components at (3). **b**, Users can search by human gene name, synonym, Entrez gene ID or UniProt accession (1). Searches are compatible with regular expressions. All records matching the search term will be reported (2), with a list of viral baits that detected the protein at an FDR ≤ 0.01. The “Profile” link (3) will display a heat map profile of the protein across all viral baits. **c**, After clicking on a “Profile” link, a heat map will be displayed of all significant proximity interactions. All preys detected with an FDR ≤ 0.01 are present and both baits and preys are clustered. Search for a prey or view the legend from (1). The heat map (2) will be scrolled to focus on a search result or when linking from an entry on the search page as shown in **b**.

**Extended Data Fig. 5.**
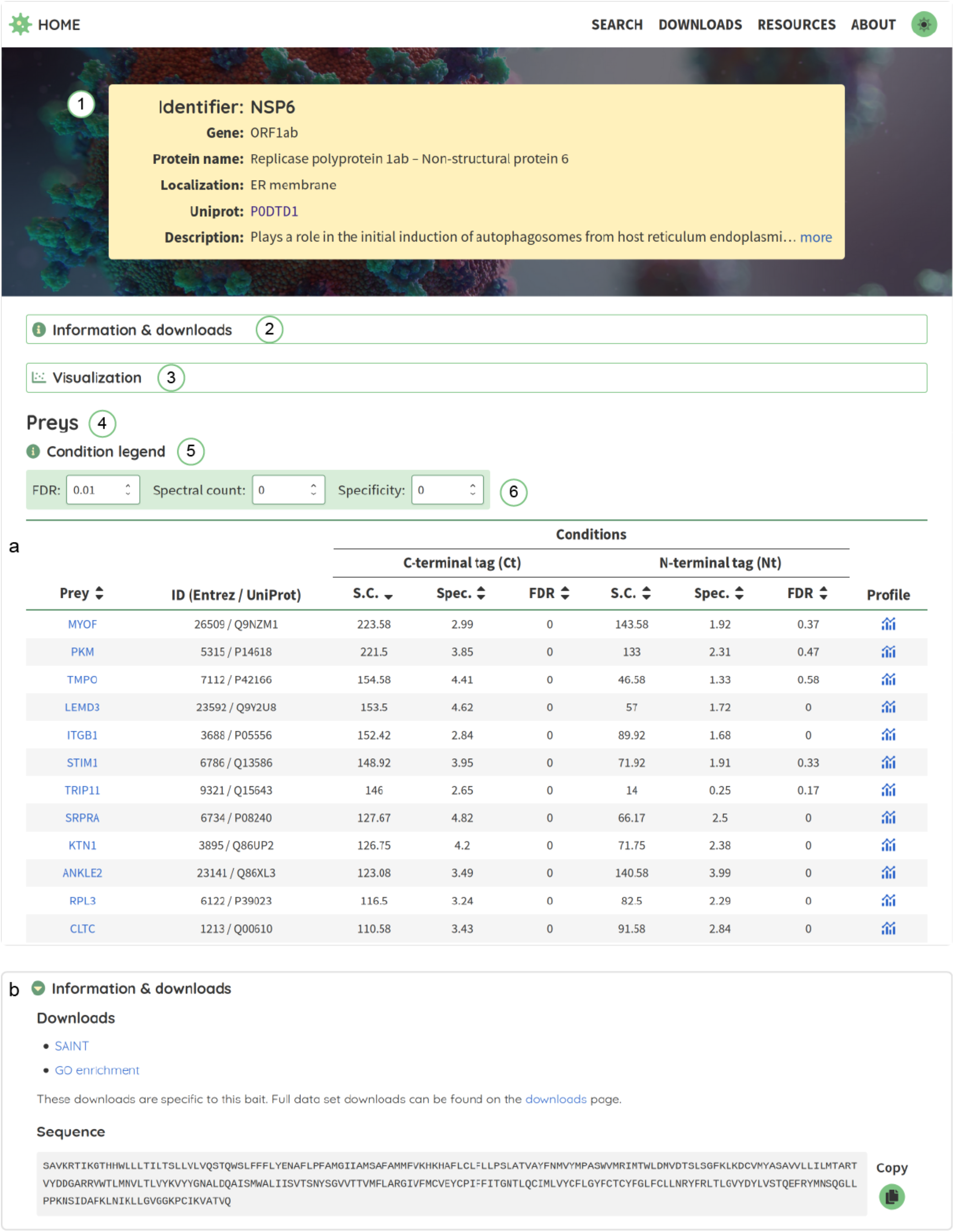
Bait report. **a**, All viral baits have a report page that begins with basic information about the protein (1). Downloads and sequence information can be found at (2) (more in panel **b**). Tools for visualizing bait results can be found in (3) (more in **Extended Data Fig. 6**). A table of preys detected with the bait occupies the bulk of the report page (4). Any conditions for the bait, including N- and C-terminal tags and mutants will be reported. The table can be sorted by prey symbol, spectral count, specificity or FDR by clicking on the respective header. Acronym definitions are found in (5) and the table can be filtered from (6). Spectral counts reported in the table are control-subtracted (see **Methods** for details). **b**, Information and data on the bait available for download can be found in the “Information & downloads” section.

**Extended Data Fig. 6.**
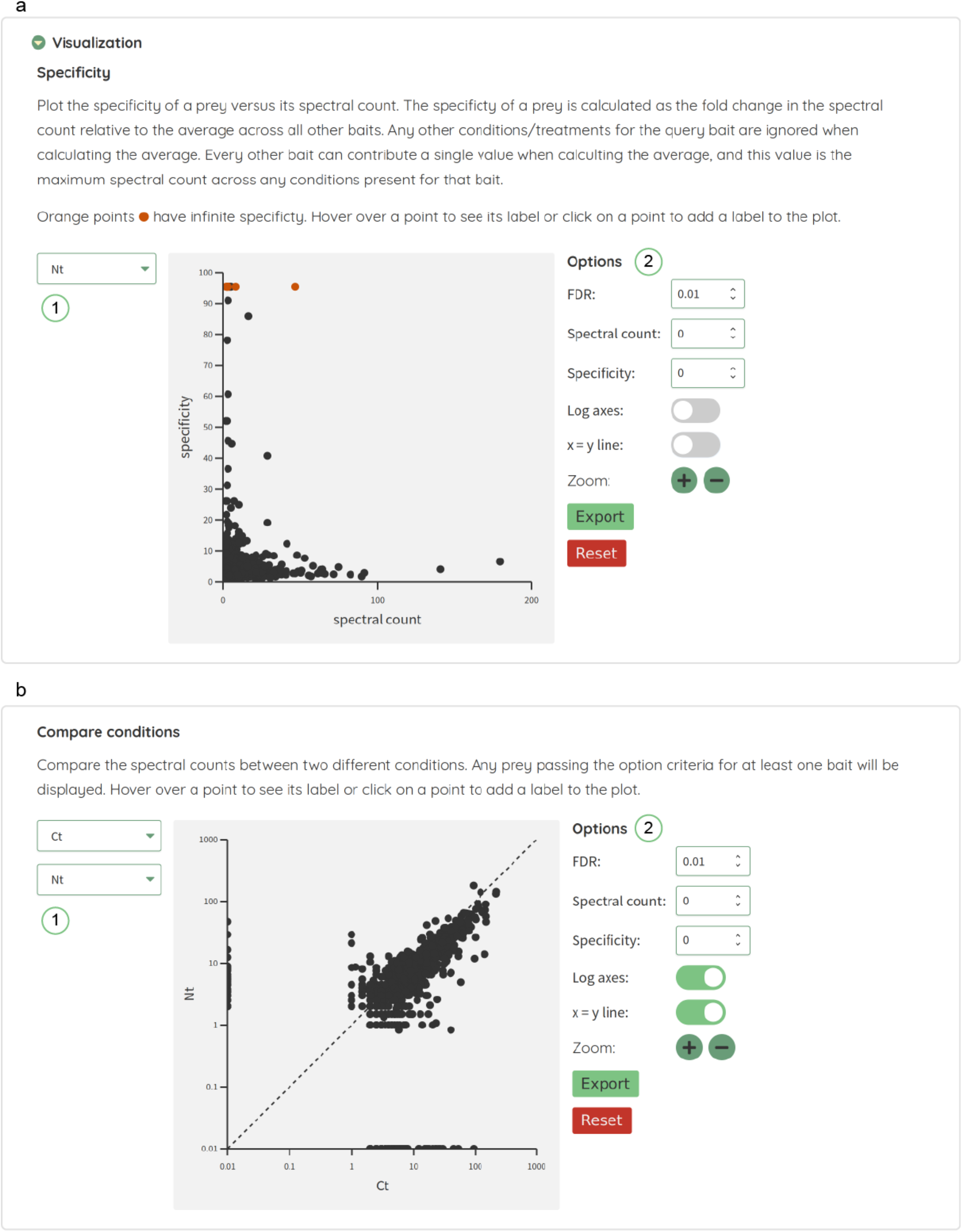
Bait visualization tools. Tools for visualizing preys graphically are available in the “Visualization” section. **a**, Specificity plots for a bait condition can be viewed by selecting the condition of interest from (1). This will display all preys that pass the filter criteria by spectral count (x-axis) and specificity (y-axis). There are various options for filtering the displayed points and customizing the image at (2). The image is interactive: click and drag to pan the image and use the mouse wheel (or pinch on mobile) to zoom. **b**, Condition comparison plots can be viewed by selecting two conditions from (1). This will display all preys that pass the filter criteria for at least one bait. Preys will be displayed by spectral count of one bait versus the other. Options for filtering and interactivity are as described for panel **a**.

**Extended Data Fig. 7.**
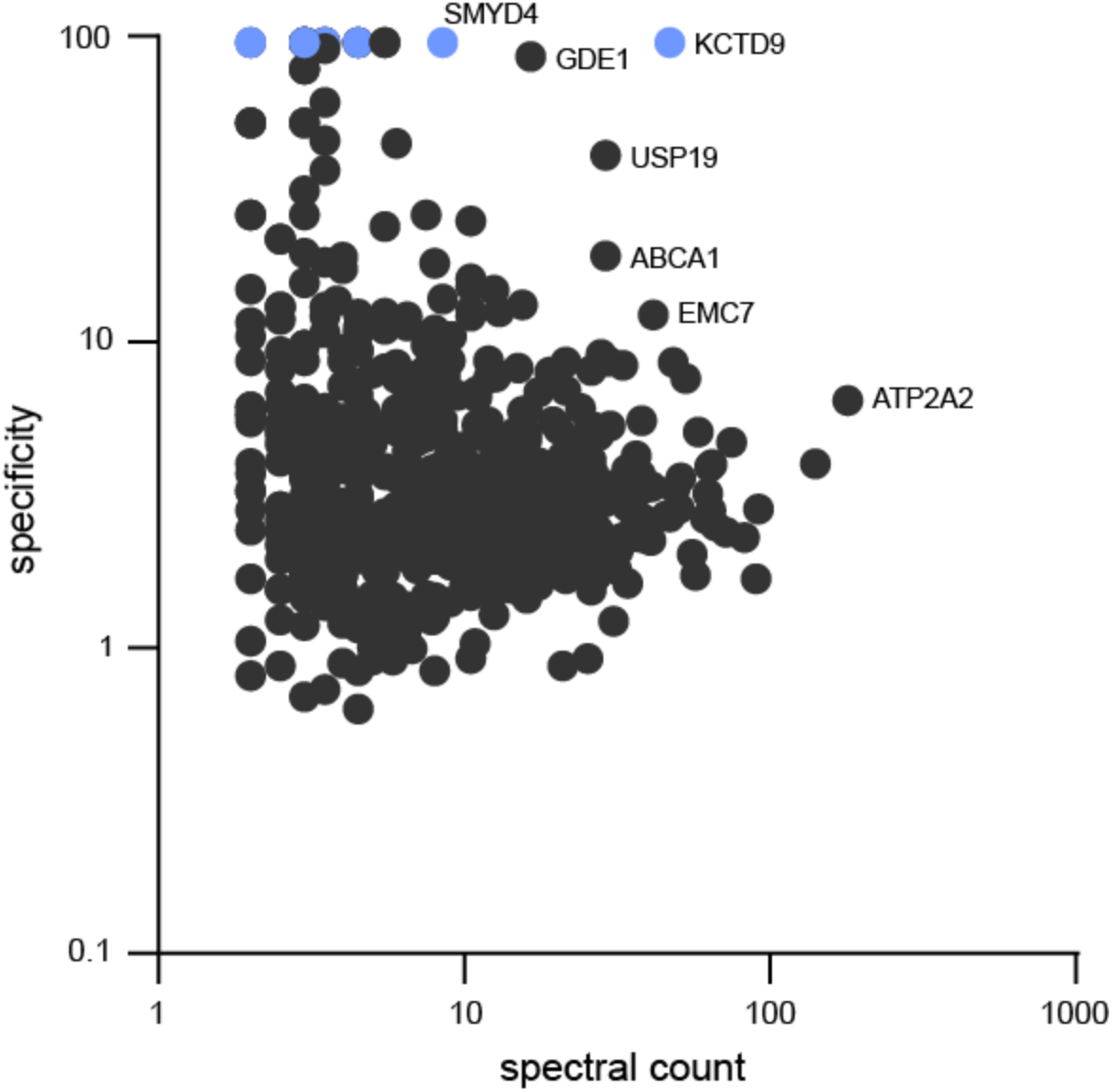
Specific interactors for NSP6_Nt (supports Fig. 3e). Spectral counts (x-axis) versus quantitative enrichment (i.e. specificity, y-axis) for preys in the NSP6_Nt BioID. Selected preys are indicated.

**Extended Data Fig. 8:**
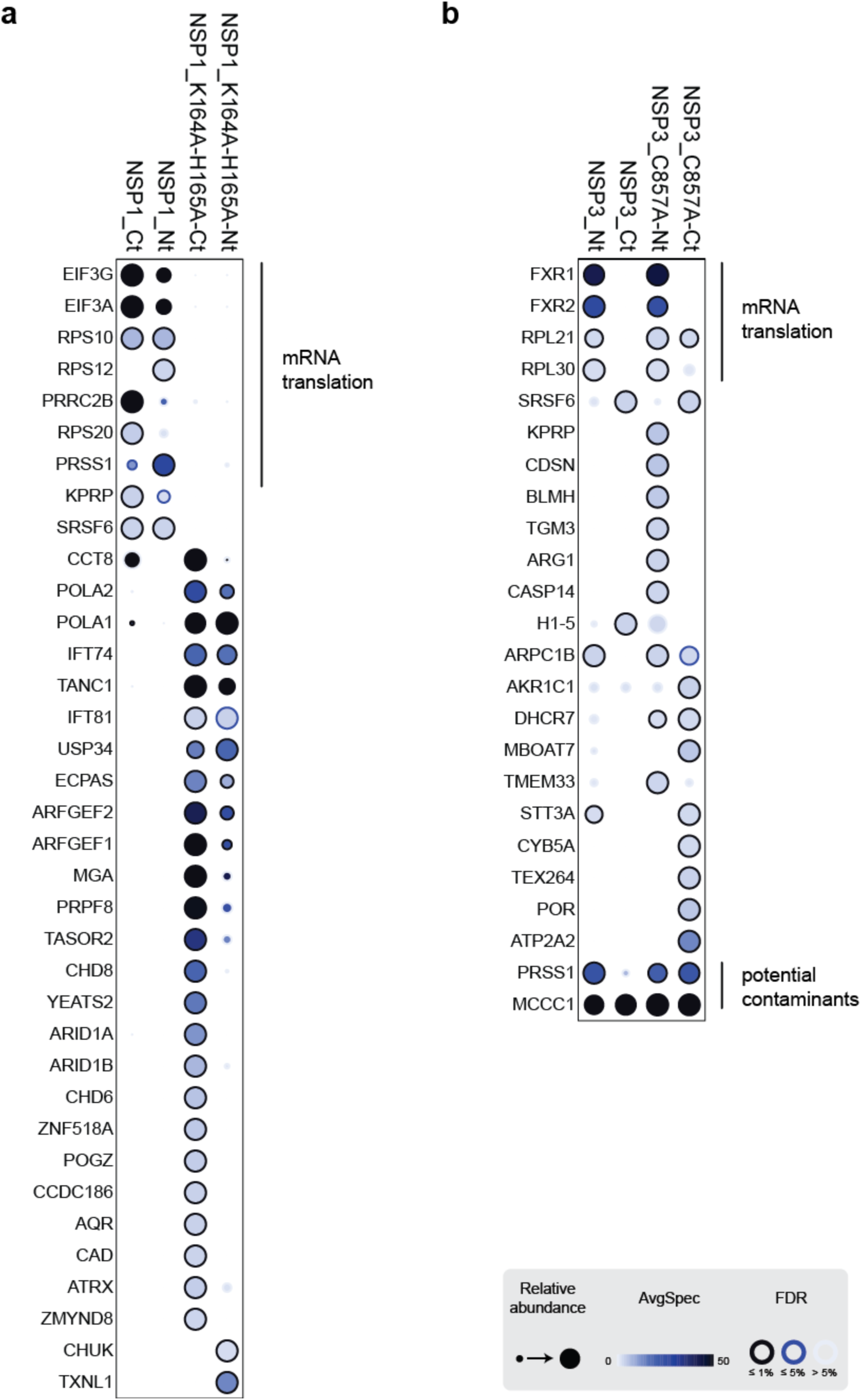
Proximity interactomes for NSP1, NSP3 wild-type and mutants (supports Fig. 4a). Dotplot view of all high-confidence proximity interactors for NSP1 and NSP3 wild type and mutant proteins. Proteins with a role in mRNA translation are indicated.

**Extended data Fig. 9:**
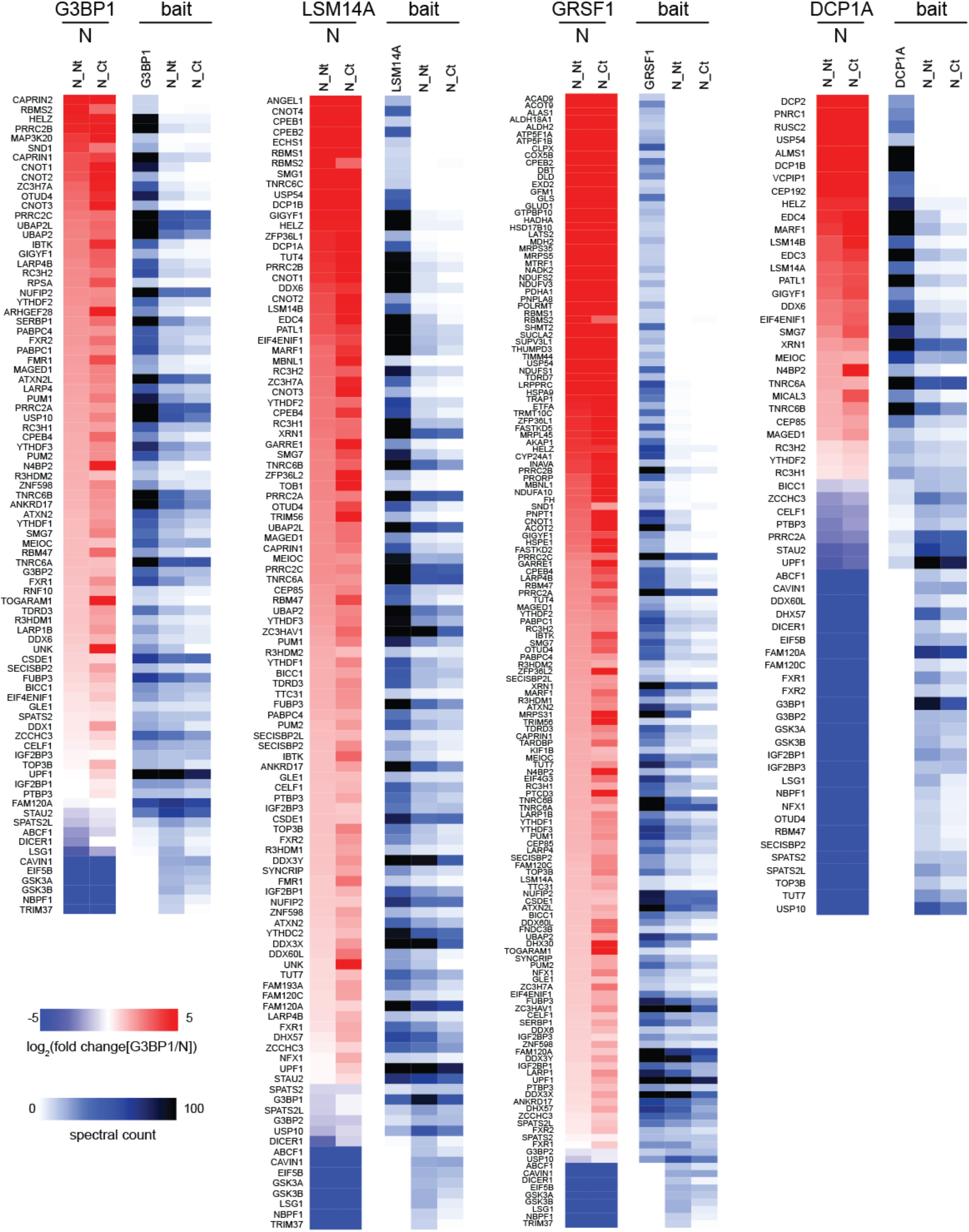
Comparison of N with RNA granule components (supports Fig. 4b). Each panel represents the prey enrichment for the indicated host RNA granule protein and N, tagged at either terminus. Two heatmaps are shown for each host bait. Left heatmap: fold change in spectral counts for preys detected with the RNA granule protein relative to N. Preys had to have an FDR of at least 0.01 and control-subtracted spectral counts of at least ten for either the RNA granule protein or N. Only preys with a consistent direction of change between N_Nt and N_Ct are shown. The fold change was capped at +/− 5 since many values are infinite (i.e. only detected with the RNA granule protein or N). Preys in the image are sorted by N_Nt and then by prey name when preys have equivalent fold changes. Right heat map: averaged spectral count for each indicated bait-prey pair.

**Extended Data Fig. 10.**
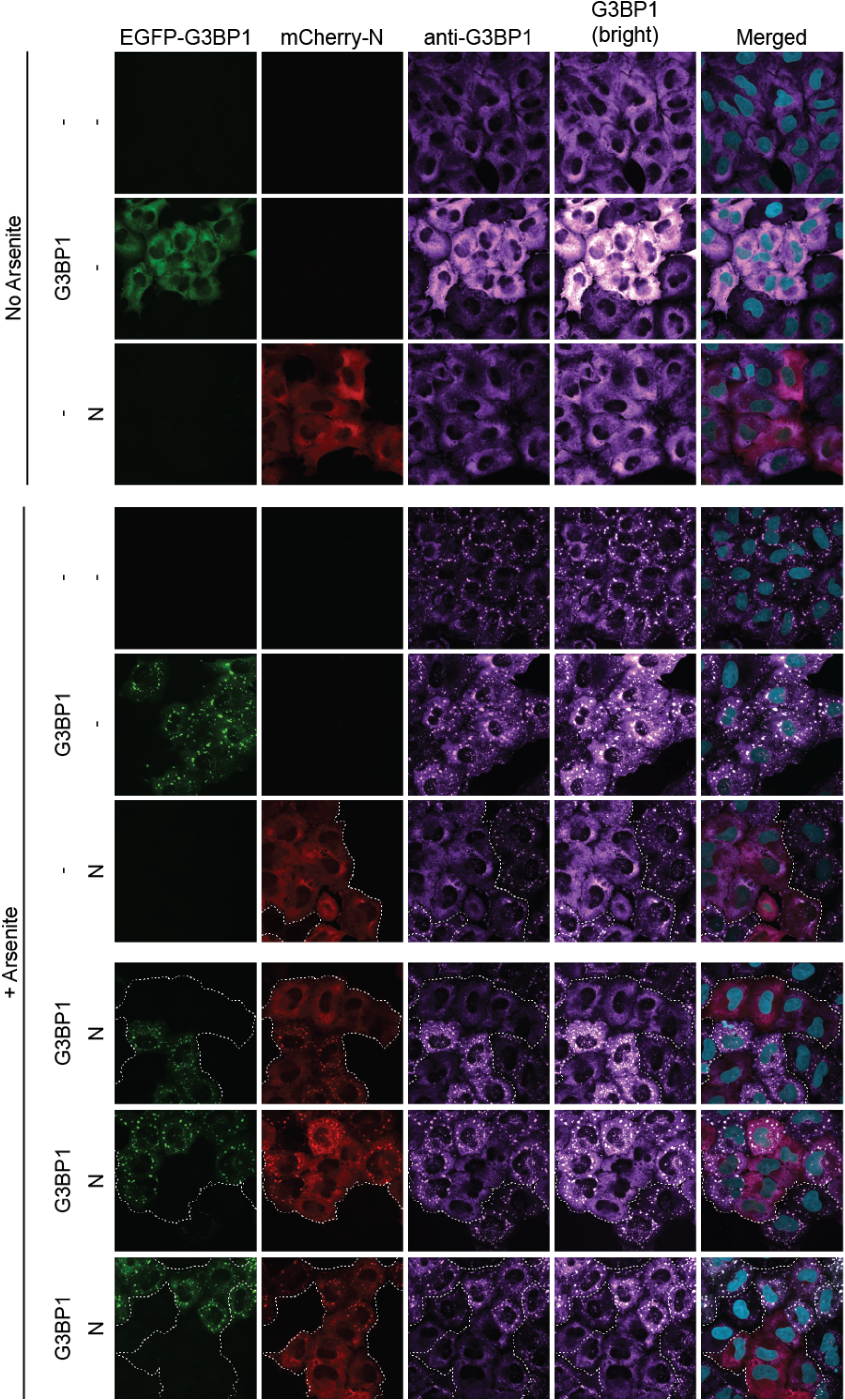
Co-localization of N in G3BP1-stress granules upon overexpression of exogenous G3BP1 (supports Fig. 4c). Using a lentivirus delivery system, mCherry-N and/or EGFP-G3BP1 were introduced into A549 cells under the control of a doxycycline inducible promoter. After induction of expression for 24 hourscells were treated with 0.5 mM sodium arsenite for 30 minutes prior to fixing with 4% paraformaldehyde. Cells were processed for immunofluorescence as described in the Methods section, and stained for endogenous anti-G3BP1, detected using Alexa647-conjugated secondary antibody, and imaged using confocal microscopy. For G3BP1/N transduced cells (bottom three panels), three independent fields are shown.

## References

1 Wu, F. et al. A new coronavirus associated with human respiratory disease in China. Nature 579, 265–269, doi:10.1038/s41586-020-2008-3 (2020).

2 Kim, D. et al. The Architecture of SARS-CoV-2 Transcriptome. Cell 181, 914–921.e910, doi:10.1016/j.cell.2020.04.011 (2020).

3 Chan, J. F. et al. Genomic characterization of the 2019 novel human-pathogenic coronavirus isolated from a patient with atypical pneumonia after visiting Wuhan. Emerg Microbes Infect 9, 221–236, doi:10.1080/22221751.2020.1719902 (2020).

4 Belov, G. A. & van Kuppeveld, F. J. (+)RNA viruses rewire cellular pathways to build replication organelles.

5 Blanchard, E. & Roingeard, P. Virus-induced double-membrane vesicles. Cellular microbiology 17, 45–50, doi:10.1111/cmi.12372 (2015).

6 Gordon, D. E. et al. A SARS-CoV-2 protein interaction map reveals targets for drug repurposing. Nature 583, 459–468, doi:10.1038/s41586-020-2286-9 (2020).

7 Stukalov, A. et al. Multi-level proteomics reveals host-perturbation strategies of SARS-CoV-2 and SARS-CoV. bioRxiv, 2020.2006.2017.156455, doi:10.1101/2020.06.17.156455 (2020).

8 Li, J. et al. Virus-host interactome and proteomic survey of PMBCs from COVID-19 patients reveal potential virulence factors influencing SARS-CoV-2 pathogenesis. bioRxiv, 2020.2003.2031.019216, doi:10.1101/2020.03.31.019216 (2020).

9 Gingras, A. C., Abe, K. T. & Raught, B. Getting to know the neighborhood: using proximity-dependent biotinylation to characterize protein complexes and map organelles. Curr Opin Chem Biol 48, 44–54, doi:10.1016/j.cbpa.2018.10.017 (2019).

10 Roux, K. J., Kim, D. I., Raida, M. & Burke, B. A promiscuous biotin ligase fusion protein identifies proximal and interacting proteins in mammalian cells. J Cell Biol 196, 801–810, doi:10.1083/jcb.201112098 (2012).

11 Couzens, A. L. et al. Protein interaction network of the mammalian Hippo pathway reveals mechanisms of kinase-phosphatase interactions. Sci Signal 6, rs15, doi:10.1126/scisignal.2004712 (2013).

12 St-Denis, N. et al. Phenotypic and Interaction Profiling of the Human Phosphatases Identifies Diverse Mitotic Regulators. Cell Rep 17, 2488–2501, doi:10.1016/j.celrep.2016.10.078 (2016).

13 Youn, J. Y. et al. High-Density Proximity Mapping Reveals the Subcellular Organization of mRNA-Associated Granules and Bodies. Mol Cell 69, 517–532 e511, doi:10.1016/j.molcel.2017.12.020 (2018).

14 Gupta, G. D. et al. A Dynamic Protein Interaction Landscape of the Human Centrosome-Cilium Interface. Cell 163, 1484–1499, doi:10.1016/j.cell.2015.10.065 (2015).

15 Coyaud, E. et al. Global Interactomics Uncovers Extensive Organellar Targeting by Zika Virus. Mol Cell Proteomics 17, 2242–2255, doi:10.1074/mcp.TIR118.000800 (2018).

16 Samavarchi-Tehrani, P., Samson, R. & Gingras, A. C. Proximity Dependent Biotinylation: Key Enzymes and Adaptation to Proteomics Approaches. Mol Cell Proteomics 19, 757–773, doi:10.1074/mcp.R120.001941 (2020).

17 Branon, T. C. et al. Efficient proximity labeling in living cells and organisms with TurboID. Nat Biotechnol 36, 880–887, doi:10.1038/nbt.4201 (2018).

18 Samavarchi-Tehrani, P., Abdouni, H., Samson, R. & Gingras, A. C. A Versatile Lentiviral Delivery Toolkit for Proximity-dependent Biotinylation in Diverse Cell Types. Mol Cell Proteomics 17, 2256–2269, doi:10.1074/mcp.TIR118.000902 (2018).

19 Kim, D. K. et al. A Comprehensive, Flexible Collection of SARS-CoV-2 Coding Regions. G3 (Bethesda), doi:10.1534/g3.120.401554 (2020).

20 Go, C. D. et al. A proximity biotinylation map of a human cell. bioRxiv, 796391, doi:10.1101/796391 (2019).

21 Kamitani, W., Huang, C., Narayanan, K., Lokugamage, K. G. & Makino, S. A two-pronged strategy to suppress host protein synthesis by SARS coronavirus Nsp1 protein. Nat Struct Mol Biol 16, 1134–1140, doi:10.1038/nsmb.1680 (2009).

22 Narayanan, K. et al. Severe acute respiratory syndrome coronavirus nsp1 suppresses host gene expression, including that of type I interferon, in infected cells. J Virol 82, 4471–4479, doi:10.1128/JVI.02472-07 (2008).

23 Teo, G. et al. SAINTexpress: improvements and additional features in Significance Analysis of INTeractome software. J Proteomics 100, 37–43, doi:10.1016/j.jprot.2013.10.023 (2014).

24 Oughtred, R. et al. The BioGRID interaction database: 2019 update. Nucleic Acids Res 47, D529–D541, doi:10.1093/nar/gky1079 (2019).

25 Dominguez Andres, A. et al. SARS-CoV-2 ORF9c Is a Membrane-Associated Protein that Suppresses Antiviral Responses in Cells. bioRxiv, 2020.2008.2018.256776, doi:10.1101/2020.08.18.256776 (2020).

26 Zhang, Y. et al. The ORF8 Protein of SARS-CoV-2 Mediates Immune Evasion through Potently Downregulating MHC-I. bioRxiv, 2020.2005.2024.111823, doi:10.1101/2020.05.24.111823 (2020).

27 te Velthuis, A. J., Arnold, J. J., Cameron, C. E., van den Worm, S. H. & Snijder, E. J. The RNA polymerase activity of SARS-coronavirus nsp12 is primer dependent. Nucleic Acids Res 38, 203–214, doi:10.1093/nar/gkp904 (2010).

28 Imbert, I. et al. A second, non-canonical RNA-dependent RNA polymerase in SARS coronavirus. EMBO J 25, 4933–4942, doi:10.1038/sj.emboj.7601368 (2006).

29 Angelini, M. M., Akhlaghpour, M., Neuman, B. W. & Buchmeier, M. J. Severe acute respiratory syndrome coronavirus nonstructural proteins 3, 4, and 6 induce double-membrane vesicles. mBio 4, doi:10.1128/mBio.00524-13 (2013).

30 Clementz, M. A., Kanjanahaluethai, A., O’Brien, T. E. & Baker, S. C. Mutation in murine coronavirus replication protein nsp4 alters assembly of double membrane vesicles. Virology 375, 118–129, doi:10.1016/j.virol.2008.01.018 (2008).

31 Di Mattia, T. et al. Identification of MOSPD2, a novel scaffold for endoplasmic reticulum membrane contact sites. EMBO Rep 19, doi:10.15252/embr.201745453 (2018).

32 Jiang, H. W. et al. SARS-CoV-2 Orf9b suppresses type I interferon responses by targeting TOM70. Cell Mol Immunol, doi:10.1038/s41423-020-0514-8 (2020).

33 Ren, Z. et al. Regulation of MAVS Expression and Signaling Function in the Antiviral Innate Immune Response. Frontiers in immunology 11, 1030, doi:10.3389/fimmu.2020.01030 (2020).

34 Antonicka, H. et al. A High-Density Human Mitochondrial Proximity Interaction Network. Cell Metabolism 32, 479–497.e479, doi:https://doi.org/10.1016/j.cmet.2020.07.017 (2020).

35 Han, L. et al. SARS-CoV-2 ORF9b Antagonizes Type I and III Interferons by Targeting Multiple Components of RIG-I/MDA-5-MAVS, TLR3-TRIF, and cGAS-STING Signaling Pathways. bioRxiv, 2020.2008.2016.252973, doi:10.1101/2020.08.16.252973 (2020).

36 Jaafar, Z. A. & Kieft, J. S. Viral RNA structure-based strategies to manipulate translation. Nature reviews. Microbiology 17, 110–123, doi:10.1038/s41579-018-0117-x (2019).

37 Thoms, M. et al. Structural basis for translational shutdown and immune evasion by the Nsp1 protein of SARS-CoV-2. Science, eabc8665, doi:10.1126/science.abc8665 (2020).

38 Chiu, W. L. et al. The C-terminal region of eukaryotic translation initiation factor 3a (eIF3a) promotes mRNA recruitment, scanning, and, together with eIF3j and the eIF3b RNA recognition motif, selection of AUG start codons. Mol Cell Biol 30, 4415–4434, doi:10.1128/mcb.00280-10 (2010).

39 Aitken, C. E. et al. Eukaryotic translation initiation factor 3 plays distinct roles at the mRNA entry and exit channels of the ribosomal preinitiation complex. Elife 5, doi:10.7554/eLife.20934 (2016).

40 Chaudhuri, A. Comparative analysis of non structural protein 1 of SARS-COV2 with SARS-COV1 and MERS-COV: An i*n silico* study. bioRxiv, 2020.2006.2009.142570, doi:10.1101/2020.06.09.142570 (2020).

41 Egloff, M.-P. et al. The severe acute respiratory syndrome-coronavirus replicative protein nsp9 is a single-stranded RNA-binding subunit unique in the RNA virus world. Proceedings of the National Academy of Sciences of the United States of America 101, 3792–3796, doi:10.1073/pnas.0307877101 (2004).

42 Li, Y. et al. LSm14A is a processing body-associated sensor of viral nucleic acids that initiates cellular antiviral response in the early phase of viral infection. Proceedings of the National Academy of Sciences 109, 11770–11775, doi:10.1073/pnas.1203405109 (2012).

43 Brandmann, T. et al. Molecular architecture of LSM14 interactions involved in the assembly of mRNA silencing complexes. EMBO J 37, doi:10.15252/embj.201797869 (2018).

44 Nishimura, T. et al. The eIF4E-Binding Protein 4E-T Is a Component of the mRNA Decay Machinery that Bridges the 5′ and 3′ Termini of Target mRNAs. Cell Reports 11, 1425–1436, doi:https://doi.org/10.1016/j.celrep.2015.04.065 (2015).

45 Hein, M. Y. et al. A human interactome in three quantitative dimensions organized by stoichiometries and abundances.

46 Huttlin, E. L. et al. Architecture of the human interactome defines protein communities and disease networks.

47 Jain, D. et al. ketu mutant mice uncover an essential meiotic function for the ancient RNA helicase YTHDC2. eLife 7, e30919, doi:10.7554/eLife.30919 (2018).

48 Gosselin, P. et al. Tracking a refined eIF4E-binding motif reveals Angel1 as a new partner of eIF4E.

49 Youn, J. Y. et al. Properties of Stress Granule and P-Body Proteomes. Mol Cell 76, 286–294, doi:10.1016/j.molcel.2019.09.014 (2019).

50 Yen, Y. T. et al. Modeling the early events of severe acute respiratory syndrome coronavirus infection in vitro.

51 Corman, V. A.-O. et al. Link of a ubiquitous human coronavirus to dromedary camels.

52 Chan, J. F. et al. Differential cell line susceptibility to the emerging novel human betacoronavirus 2c EMC/2012: implications for disease pathogenesis and clinical manifestation.

53 Miller, C. L. Stress Granules and Virus Replication.

54 Perdikari, T. M. et al. SARS-CoV-2 nucleocapsid protein undergoes liquid-liquid phase separation stimulated by RNA and partitions into phases of human ribonucleoproteins. LID – 2020.06.09.141101 [pii] LID – 10.1101/2020.06.09.141101 [doi].

55 Cascarina, S. M. & Ross, E. D. A proposed role for the SARS-CoV-2 nucleocapsid protein in the formation and regulation of biomolecular condensates. LID – 10.1096/fj.202001351 [doi].

56 Savastano, A., de Opakua, A. I., Rankovic, M. & Zweckstetter, M. Nucleocapsid protein of SARS-CoV-2 phase separates into RNA-rich polymerase-containing condensates. bioRxiv, 2020.2006.2018.160648, doi:10.1101/2020.06.18.160648 (2020).

57 Iserman, C. et al. Specific viral RNA drives the SARS CoV-2 nucleocapsid to phase separate. bioRxiv, 2020.2006.2011.147199, doi:10.1101/2020.06.11.147199 (2020).

58 Cho, K. F. et al. Split-TurboID enables contact-dependent proximity labeling in cells. Proc Natl Acad Sci U S A 117, 12143–12154, doi:10.1073/pnas.1919528117 (2020).

59 Hoffmann, M. et al. SARS-CoV-2 Cell Entry Depends on ACE2 and TMPRSS2 and Is Blocked by a Clinically Proven Protease Inhibitor. Cell 181, 271–280.e278, doi:10.1016/j.cell.2020.02.052 (2020).

60 Hikmet, F. et al. The protein expression profile of ACE2 in human tissues. Molecular systems biology 16, e9610, doi:10.15252/msb.20209610 (2020).

61 Werion, A. et al. SARS-CoV-2 Causes a Specific Dysfunction of the Kidney Proximal Tubule. Kidney international, doi:10.1016/j.kint.2020.07.019 (2020).

62 Li, Z. et al. Caution on Kidney Dysfunctions of COVID-19 Patients. medRxiv, 2020.2002.2008.20021212, doi:10.1101/2020.02.08.20021212 (2020).

63 Cheng, Y. et al. Kidney impairment is associated with in-hospital death of COVID-19 patients. medRxiv, 2020.2002.2018.20023242, doi:10.1101/2020.02.18.20023242 (2020).

64 Zheng, Y. Y., Ma, Y. T., Zhang, J. Y. & Xie, X. COVID-19 and the cardiovascular system. Nature reviews. Cardiology 17, 259–260, doi:10.1038/s41569-020-0360-5 (2020).

65 Wang, Y. et al. SARS-CoV-2 infection of the liver directly contributes to hepatic impairment in patients with COVID-19. Journal of hepatology, doi:10.1016/j.jhep.2020.05.002 (2020).

66 Yang, X. et al. Clinical course and outcomes of critically ill patients with SARS-CoV-2 pneumonia in Wuhan, China: a single-centered, retrospective, observational study. The Lancet. Respiratory medicine 8, 475–481, doi:10.1016/s2213-2600(20)30079-5 (2020).

67 Song, P., Li, W., Xie, J., Hou, Y. & You, C. Cytokine storm induced by SARS-CoV-2. Clinica chimica acta; international journal of clinical chemistry 509, 280–287, doi:10.1016/j.cca.2020.06.017 (2020).

68 Puelles, V. G. et al. Multiorgan and Renal Tropism of SARS-CoV-2. The New England journal of medicine 383, 590–592, doi:10.1056/NEJMc2011400 (2020).

69 Liu, G. et al. ProHits: integrated software for mass spectrometry-based interaction proteomics. Nat Biotechnol 28, 1015–1017, doi:10.1038/nbt1010-1015 (2010).

70 Eng, J. K., Jahan, T. A. & Hoopmann, M. R. Comet: an open-source MS/MS sequence database search tool. Proteomics 13, 22–24, doi:10.1002/pmic.201200439 (2013).

71 Keller, A., Nesvizhskii, A. I., Kolker, E. & Aebersold, R. Empirical statistical model to estimate the accuracy of peptide identifications made by MS/MS and database search. Analytical chemistry 74, 5383–5392, doi:10.1021/ac025747h (2002).

72 Shteynberg, D. et al. iProphet: multi-level integrative analysis of shotgun proteomic data improves peptide and protein identification rates and error estimates. Mol Cell Proteomics 10, M111.007690, doi:10.1074/mcp.M111.007690 (2011).

73 Mellacheruvu, D. et al. The CRAPome: a contaminant repository for affinity purification-mass spectrometry data. Nat Methods 10, 730–736, doi:10.1038/nmeth.2557 (2013).

74 Raudvere, U. et al. g:Profiler: a web server for functional enrichment analysis and conversions of gene lists (2019 update). Nucleic Acids Res 47, W191–w198, doi:10.1093/nar/gkz369 (2019).

75 Samaras, P. et al. ProteomicsDB: a multi-omics and multi-organism resource for life science research. Nucleic Acids Res 48, D1153–d1163, doi:10.1093/nar/gkz974 (2020).

76 El-Gebali, S. et al. The Pfam protein families database in 2019. Nucleic Acids Res 47, D427–d432, doi:10.1093/nar/gky995 (2019).

77 Stark, C. et al. BioGRID: a general repository for interaction datasets. Nucleic Acids Res 34, D535–539, doi:10.1093/nar/gkj109 (2006).

78 Knight, J. D. R. et al. ProHits-viz: a suite of web tools for visualizing interaction proteomics data. Nat Methods 14, 645–646, doi:10.1038/nmeth.4330 (2017).

79 Shannon, P. et al. Cytoscape: a software environment for integrated models of biomolecular interaction networks. Genome research 13, 2498–2504, doi:10.1101/gr.1239303 (2003).

